# MVCBench: A Multimodal Benchmark for Drug-induced Virtual Cell Phenotypes

**DOI:** 10.64898/2026.04.22.720110

**Authors:** Bo Li, Qing Wang, Shihang Wang, Bob Zhang, Yuzhong Peng, Pinxian Zeng, Chengliang Liu, Mengran Li, Ziyang Tang, Xiaojun Yao, Chuxia Deng, Qianqian Song

**Author notes:** Corresponding authors: Qianqian Song, PhD; Bob Zhang, PhD. The authors contributed equally to this work (Bo Li, Qing Wang, Shihang Wang).

## Abstract

Drugs induce coordinated phenotypic changes across multiple modalities, including transcriptional reprogramming and cellular morphological remodeling. Predicting these drug-induced modality changes is central to drug discovery, mechanism-of-action studies and precision therapeutics, however, prediction performance depends critically on how both drug compounds and cellular states are represented. Despite rapid advances in drug molecular and gene representation methods, a systematic evaluation of these methods remains lacking. Herein, we introduce MVCBench, a comprehensive benchmarking framework for evaluating drug molecular and gene representation methods in predicting drug-induced multimodal virtual cell (MVC) phenotypes. MVCBench leverages large-scale transcriptomic and high-content imaging data and systematically evaluates 24 representation methods (12 drug molecular and 12 gene representation methods) across nearly 1.1 million drug-induced profiles, under both in-distribution and out-of-distribution settings spanning unseen compounds, cell lines, assay plates and datasets. Our benchmarking reveals a pronounced modality-dependent asymmetry: advanced drug molecular representations substantially improve the prediction of drug-induced morphological phenotypes but provide only limited gains for gene expression prediction relative to classical fingerprints, whereas task-specific gene representations outperform general-purpose foundation models in predicting drug-induced transcriptomic responses. Predictive performance also deteriorates sharply under distribution shift, highlighting persistent challenges in cross-dataset and cross-platform generalization. We further show that integrating transcriptomic and morphological modalities consistently improves prediction accuracy, and derive practical design principles for MVC architectures, including modality-aware loss calibration and fusion strategies. Together, MVCBench provides a systematic foundation for evaluating representation methods and offers guidance for developing robust MVC models of drug-induced cellular responses.

## INTRODUCTION

Drug treatments induce complex and coordinated phenotypic changes across multiple modalities, reshaping gene expression programs, cellular morphology, and higher-order tissue organization^1–3^. These changes reflect the combined effects of drug-target interactions, downstream signaling cascades, and adaptive cellular responses, which together determine therapeutic efficacy, toxicity, and resistance^4,5^. Accurately predicting post-treatment virtual cell phenotypes is therefore central to drug discovery, mechanism-of-action studies, and the development of precision therapeutics that tailor treatment to specific cellular and patient contexts^6–9^. Recent advances in high-throughput profiling technologies have transformed our ability to measure these phenotypic responses at scale^10,11^. Large-scale transcriptomic assays enable comprehensive characterization of drug-induced regulatory and transcriptional programs^12^, while high-content cellular imaging captures rich morphological changes that reflect alterations in cell state and function^13^. Together, these modalities provide complementary and mechanistically informative views of drug-induced phenotypic changes^2,14^. The growing availability of paired pre- and post-treatment data across transcriptomic and morphological modalities has created unprecedented opportunities to predict drug-induced virtual cell phenotypes, through predictive modeling that infers post-treatment phenotypes from baseline cellular states and drug compound properties^15–17^.

Predicting drug-induced virtual cell phenotypes, such as transcriptomic and morphological responses, requires effective representations of both the drug compound and the pre-treatment biological state^18–20^. Accordingly, two complementary classes of representation methods serve as the backbone of these predictive frameworks. The first category comprises drug molecular representation methods, which transform the chemical structures of drug compounds into latent features suitable for phenotypic prediction. This category spans a diverse spectrum, ranging from rule-based explicit fingerprints (e.g., ECFP4^19^) and graph neural networks^21^ (GNN) (e.g., Chemprop^22^) to large-scale molecular foundation models (e.g., KPGT^23^). The second category comprises gene representation methods, increasingly instantiated as single-cell foundation models (scFMs). These methods capture the pre-treatment biological state by mapping high-dimensional transcriptomic profiles into compact embeddings. Gene representation methods have evolved from classical dimensionality-reduction techniques to deep, foundation-scale models (e.g., scGPT^18^) trained on millions of transcriptomic profiles, aiming to extract biologically meaningful structures that generalize across diverse conditions. Together, these drug molecular and gene representation methods define the representational backbone used to predict drug-induced virtual cell phenotypes.

Crucially, drug-induced phenotypic prediction is inherently conditional and context-dependent: the post-treatment virtual cell state arises from the interplay between the drug compound and the baseline biological context^24–27^. Even the same compound can elicit markedly different transcriptomic and morphological responses depending on cell type, baseline gene programs, or cellular states^28,29^. This dependency creates a fundamental evaluation challenge: predictive performance cannot be uniquely attributed to a specific drug or gene representation method unless the complementary input is carefully controlled to avoid confounding effects. Therefore, rigorous benchmarking requires experimental designs that disentangle these two contributions. Specifically, evaluating drug molecular representation methods requires fixing the pre-treatment gene expression input, so that performance differences reflect only the quality of drug features. Meanwhile, assessing gene representation methods requires holding the drug molecular input constant. Furthermore, this challenge is amplified in multimodal setting: a molecular representation method optimized for transcriptomic prediction may fail to generalize to morphological phenotype prediction^2,29^. Together, these considerations highlight the need for a systematic, modality-aware benchmarking framework spanning transcriptomic, morphological and multimodal settings to determine when specific representation methods succeed or fail.

Motivated by these considerations, we introduce MVCBench, a systematic benchmarking framework designed to explicitly disentangle the contributions of drug molecular representations and gene representations in predicting drug-induced multimodal virtual cell phenotypes, and to identify optimal representation methods. MVCBench adopts a progressive evaluation protocol comprising two paradigms: single task prediction (STP) and multimodal virtual cell modeling (MVC). In the STP paradigm, the model predicts a single post-treatment phenotype, enabling controlled evaluation of representation methods under two settings: (1) STP-Drug representations focus, which benchmarks drug molecular representations by predicting post-treatment gene expression or morphological phenotypes, while keeping the pre-treatment cellular measurements (gene expression or morphology) fixed. (2) STP-Gene representations focus, which benchmarks gene representation methods for predicting post-treatment gene expression with drug inputs held constant. In the MVC paradigm, drug representations are integrated with both pre-drug transcriptomic and morphological modalities to jointly predict drug-induced modalities. This setting evaluates how effectively chemical and biological information can be combined to model complex drug-induced cellular responses. By systematically alternating representation methods across these paradigms, MVCBench provides a unified and interpretable view of how drug and gene representations perform across distinct phenotypic domains. Our results reveal modality-specific strengths and limitations among widely used methods, offering practical guidance for selecting representation methods in drug-induced multimodal virtual cell modeling.

## RESULTS

### The MVCBench framework

Drug-induced phenotypic changes exhibit multimodal cellular responses, encompassing both transcriptomic reprogramming and morphological remodeling (**Fig. 1a**). To systematically evaluate the performance of drug molecular representation methods and gene representation methods in this context, we compiled a comprehensive dataset spanning two major modalities (transcriptomic, morphology) within the MVCBench framework (**Fig. 1b**). For the transcriptomic modality, five datasets were included: LINCS2020^13,30^, Tahoe_mini^31^, CIGS^32^, as well as the gene expression data (denoted as BBBC047_Gene and BBBC036_Gene) from CDRPBIO-BBBC036-Bray and CDRP-BBBC047-Bray^10^. Collectively, these datasets encompass 38,950 compounds and comprise 482,323 paired pre- and post-treatment transcriptomic profiles. For the morphology modality, we included cpg0016^33^, the largest available Cell Painting dataset, alongside the cell morphology data (denoted as BBBC047_Image and BBBC036_Image) from CDRPBIO-BBBC036-Bray and CDRP-BBBC047-Bray^10^, covering 84,858 compounds and 601,412 morphological profiles with pre- and post-treatment samples. To illustrate the scale and composition of these resources, **Supplementary Fig. 1a** shows the distribution of unique compounds and paired profiles across two modalities. In addition, inter-plate and cross-dataset heterogeneity was quantified to characterize dataset variability (**Supplementary Note 1, Supplementary Fig. 1b-c**). Detailed data information is provided in the **Benchmarking Datasets** section and **Table 1**.

**Fig. 1:**
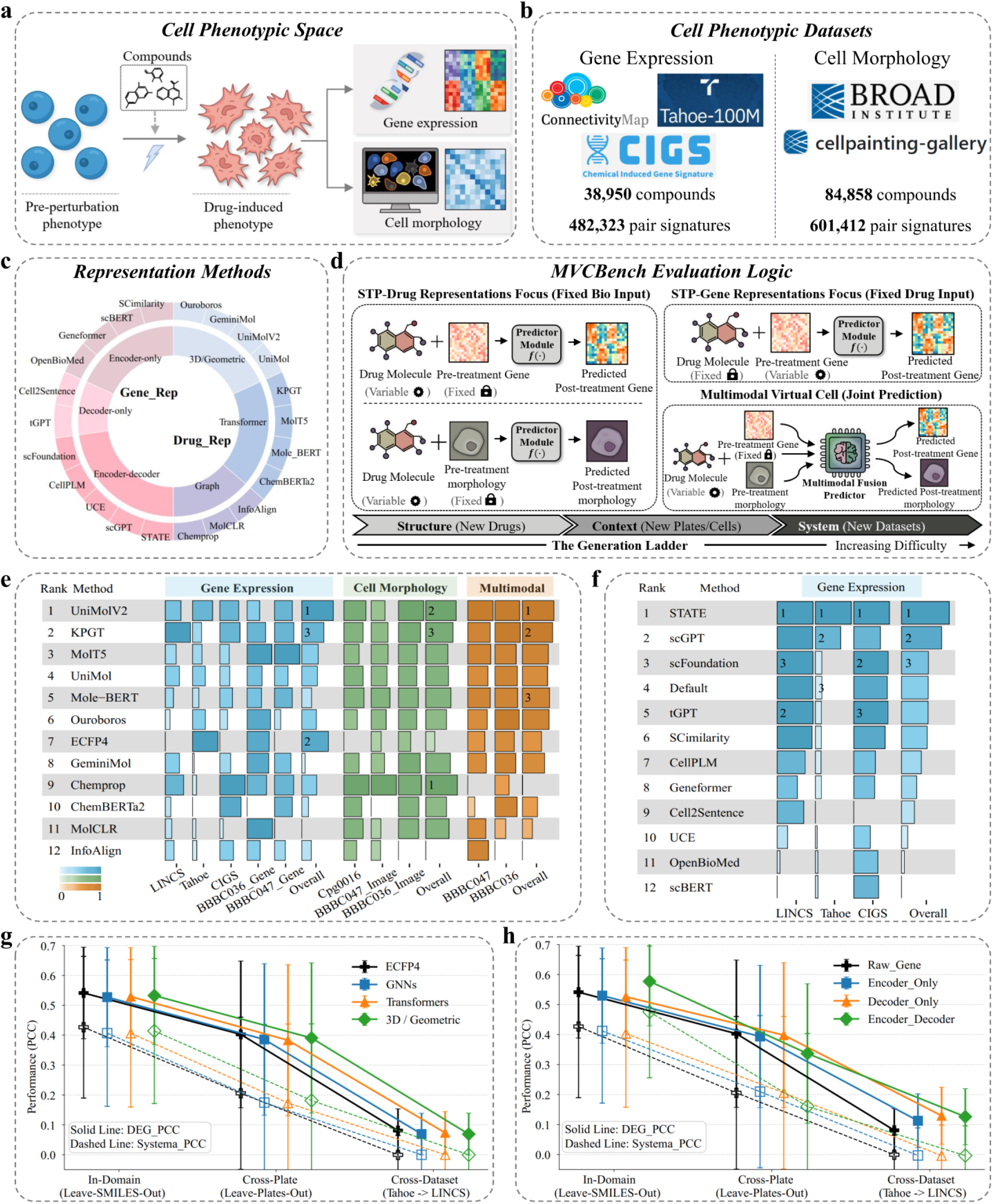
Overview of MVCBench datasets, evaluation framework, and benchmark tasks. **a**, Schematic of multimodal cellular responses, illustrating drug-induced transcriptomic and morphological responses. **b**, Composition of MVCBench datasets, integrating five transcriptomic resources (LINCS2020, Tahoe_mini, CIGS, BBBC047_Gene, and BBBC036_Gene) and three morphology datasets (cpg0016, BBBC036_Image, BBBC047_Image), covering 38,950 compounds with 482,323 paired transcriptome profiles, and 84,858 compounds with 601,412 paired morphological profiles. **c**, Model collection, including 22 advanced deep learning-based drug and gene representation methods. **d**, Systematic evaluation framework of MVCBench. The framework comprises three benchmarking scenarios: 1) STP-Drug Representations Focus: Benchmarking drug representation methods by fixing the pre-treatment biological context (gene expression or cell morphology) to predict post-treatment phenotypes.

The MVCBench framework provides a comprehensive evaluation of 24 representation methods, including 12 drug molecular representation methods and 12 gene representation methods (**Fig. 1c**). Here we established a canonical baseline using Extended-Connectivity Fingerprints (ECFP4^19^) for drugs, and raw gene expression (denoted as Raw_Gene) for genes. These baselines provide a reference for comparison against advanced representation methods. Beyond the ECFP4 baseline, the other 11 drug molecular representation methods fall into three categories: (i) graph neural network-based models including Chemprop^22^, MolCLR^34^, and InfoAlign^35^; (ii) Transformer-based models such as ChemBERTa2^36^, Mole-BERT^37^, MolT5^38^, and KPGT^23^; and (iii) models incorporating three-dimensional or geometric information, including UniMol^39^, UniMolV2^40^, GeminiMol^41^, and Ouroboros^42^. Similarly, in addition to the Raw_Gene baseline, the other 11 deep learning-based gene representation methods also span three major categories: encoder-only (Geneformer^43^, scBERT^44^, OpenBioMed^45^, SCimilarity^46^), encoder-decoder (scFoundation^47^, scGPT^18^, UCE^48^, CellPLM^49^, STATE^50^), and decoder-only (tGPT^51^, Cell2Sentence^52^). Specifications of these representation methods, including parameter size, output dimensionality, inference latency, and publication year are summarized in **Table 2**.

To enable fair and systematic comparisons of drug and gene representation methods, we adopt the unified and fully controlled MVCBench framework (**Fig. 1d**). MVCBench employs a progressive evaluation protocol that includes two paradigms, single-task paradigm (STP) and multimodal virtual cell (MVC), with three evaluation scenarios: (1) STP-Drug representations focus, in which only the drug representation is varied. Model predicts post-treatment gene expression or morphological phenotypes using fixed pre-treatment cellular measurements (e.g., baseline gene expression or cell morphology), thereby isolating the impact of molecular encoding on phenotypic prediction (**Fig. 1d**, left). (2) STP-Gene representations focus, in which drug inputs are fixed to a baseline molecular representation and only the gene representation method is varied. This setting enables direct comparison of how different gene representations influence prediction performance under identical drug molecular encodings (**Fig. 1d**, top right). (3) Multimodal virtual cell (MVC), in which drug encodings are integrated with pre-treatment transcriptomic and morphological states to jointly predict post-treatment outcomes. This evaluation determines how effectively chemical and biological information are integrated to predict complex, drug-induced cellular responses (**Fig. 1d**, bottom right).

All three MVCBench scenarios are evaluated using a hierarchical set of increasingly challenging generalization settings, including evaluation on previously unseen drug molecules (leave-SMILES-out), unseen experimental conditions such as new assay batches (leave-plate-out) or cell lines (leave-cell-line-out), and independent datasets not used during training (leave-dataset-out). To ensure consistency, the overall prediction model in each scenario is trained and evaluated under identical experimental conditions, including the same data splits, optimization protocols, hyperparameters, and random seeds. By varying only a single input representation while keeping all other components fixed, MVCBench minimizes confounding factors and ensures that observed performance differences can be directly attributed to the quality of the underlying chemical or gene representations. This controlled design enables systematic and fair benchmarking across transcriptomic, morphological, and multimodal prediction tasks.

The central findings of this study are summarized in **Fig. 1e–h**, providing a high-level overview representation performance across various tasks and evaluation settings. **Fig. 1e** and **Fig. 1f** present in-domain rankings for the 12 drug molecular and 12 gene representation methods, respectively. These results reveal a striking asymmetry in performance: compared to the ECFP4 baseline, advanced drug molecular representations (most notably UniMolV2 and KPGT) yield substantial improvements in predicting morphological phenotypes, yet offer only marginal gains for gene expression prediction. **Fig. 1g** and **Fig. 1h** highlight out-of-distribution (OOD) generalization as the major limiting factor across all methods, with performance declining sharply as the domain shift increases across plates and independent datasets. The following sections present a detailed analysis using the MVCBench framework to evaluate the contributions and limitations of each representation method in different scenarios.

2) STP-Gene Representations Focus: Benchmarking gene representation methods by fixing drug inputs. 3) Multimodal Virtual Cell: Benchmarking drug representation methods by fixing pre-treatment multimodal biological contexts (gene expression and cell morphology), while systematically evaluating multimodal fusion strategies and loss functions for joint transcriptomic and morphological prediction. **e-f**, Performance overview of drug molecular (e) and gene (f) representation methods. Overall ranking of models under the leave-SMILES-out setting, providing a direct guidance for representation method selection. **g-h**, Performance collapse on OOD tasks and metric inflation. Line plots illustrate that representation methods degrade significantly on out-of-distribution tasks.

### Benchmarking Drug Molecular Representations for Predicting Drug-Induced Gene Expression

We first conducted a systematic benchmark of 12 drug molecular representation methods for predicting drug-induced gene expression. This analysis aims to assess how different drug molecular encodings influence post-treatment transcriptomic predictions when integrated with pre-treatment cellular states (**Fig. 1d**, *STP-Drug representations focus*). Evaluation used a leave-SMILES-out strategy (see **Methods: Evaluation Protocol**), which prevents data leakage by ensuring that compounds in the test set are not seen during training, thereby simulating prediction for truly novel drugs. Prediction performance was quantified using a suite of complementary metrics, including DEG_PCC, DEG_RMSE, Systema_PCC^53^, and Systema_RMSE, with detailed definitions provided in **Methods: Evaluation Metrics** section.

**Fig. 2a** shows that different drug molecular representation methods result in marginal performance variations on the LINCS2020 dataset, with top-performing models differing only at the second decimal place. Among these drug molecular representation methods, KPGT achieved the highest median DEG_PCC (0.609), followed closely by UniMolV2 and Chemprop (0.608), while GeminiMol, InfoAlign, and similar architectures scored slightly lower (0.606). The explicit rule-based baseline (ECFP4) remained robust (0.604), showing a performance gap of less than 1% compared to most learned representations. Similar trends were observed for Systema_PCC (**Supplementary Fig. 2a**).

**Fig. 2:**
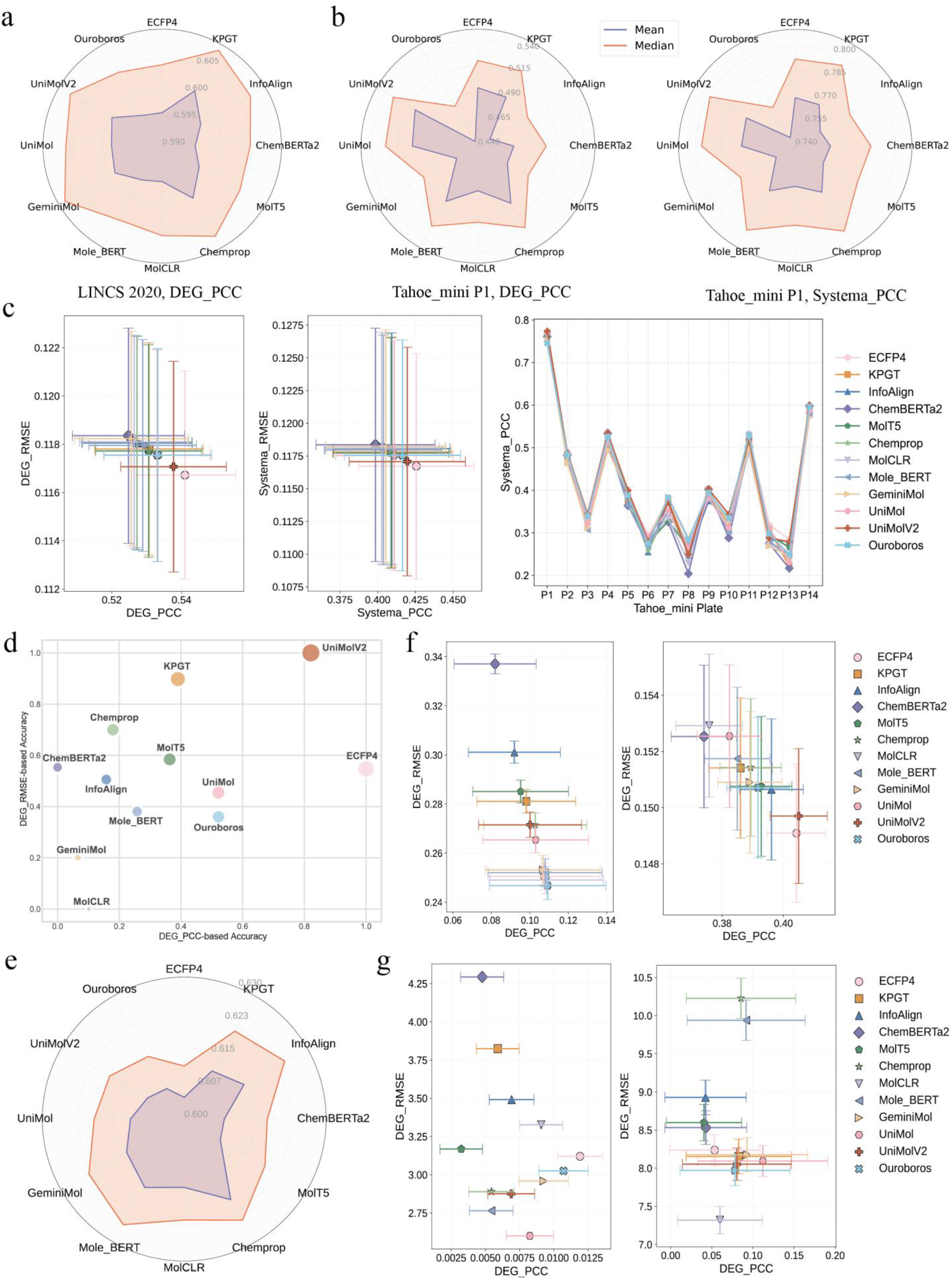
Benchmark drug molecular representation methods for drug-induced gene expression prediction. **a, b**, Performance evaluation using the leave-SMILES-out strategy on the LINCS2020 (**a**) and Tahoe_mini P1 (**b**) datasets. The inner and outer rings of the radar charts represent the mean and median values of the DEG_PCC metric, respectively. **c**, Performance summary of all 14 experimental plates in the Tahoe_mini dataset (mean ± standard deviation). From left to right: the distribution of gene-level error (DEG_RMSE), system-level correlation (Systema_PCC), and line plots of specific scores for each plate. **d**, Comprehensive performance overview across five transcriptomic datasets (LINCS2020, Tahoe_mini, CIGS, BBBC047_Gene, BBBC036_Gene). **e**, Cross-cell-line generalization test (leave-cell-line-out) on the LINCS2020 dataset. **f**, leave-plate-out generalization evaluation on the Tahoe_mini dataset. The left and right panels illustrate a significant decline in performance when using the highly heterogeneous P1 and P13 plates as the training set and the remaining plates as the test set. **g**, Bidirectional cross-dataset (leave-dataset-out) transfer tests between LINCS2020 and Tahoe_mini, demonstrating a collapse in predictive performance due to severe distribution shifts.

On the Tahoe_mini P1 dataset (**Fig. 2b**), UniMolV2, Chemprop, and Mole-BERT achieved comparable performance in Systema_PCC (medians: 0.798–0.799). ECFP4 remained competitive (median: 0.792), showing less than a 1% performance gap compared to the leading models. However, more distinct differentiation was observed in DEG_PCC, where UniMolV2 (median: 0.532) clearly outperformed the cluster of Chemprop, Mole-BERT, and UniMol. Consistent trends were observed in DEG_RMSE and Systema_RMSE (**Supplementary Fig. 2b**).

Extending this evaluation across all 14 Tahoe_mini plates (**Fig. 2c**) confirmed consistent trends with minimal performance variance among representations. The performance gap between the 12 drug molecular representation methods remained within 1%–2% (mean ± s.d.). Notably, ECFP4 achieved a DEG_PCC of 0.541 ± 0.145, while UniMolV2 (0.538 ± 0.151) and UniMol (0.533 ± 0.152) showed nearly indistinguishable performance. Chemprop and Mole-BERT performed slightly lower but remained within ±0.01 of the top methods. Additionally, the corresponding Systema_PCC metric revealed a consistent ranking pattern with the DEG_PCC results (**Fig. 2c, middle**). Furthermore, the detailed per-plate analysis (**Fig. 2c, right**) revealed that predictive fidelity is heavily plate-dependent, with Systema_PCC values varying from 0.2 (P13) to 0.6 (P14). This suggests that across different technical batches, current learned representations have not yet established a decisive advantage over classical ECFP4 fingerprints in the drug-induced gene expression prediction task. **Fig. 2d** summarizes the overall performance across all datasets (LINCS2020, Tahoe_mini, CIGS, BBBC047_Gene, and BBBC036_Gene). To provide a holistic assessment, we evaluated the models by jointly considering prediction correlation (normalized DEG_PCC) and error magnitude (normalized DEG_RMSE). The resulting distribution in the dual-metric space showed that UniMolV2 achieved the top rank (normalized DEG_PCC=0.82, normalized DEG_RMSE=1.00), overall outperforming the ECFP4 baseline (1.00, 0.55), followed by KPGT (0.39, 0.90) and MolT5 (0.36, 0.59). Other methods, such as UniMol, Chemprop, and Ouroboros, formed a secondary performance tier. Overall, most drug molecular representation methods yielded only marginal gains over the ECFP4 in gene expression prediction, with minimal performance variation across the evaluated methods.

### Generalization of Drug Molecular Representations across Cell Lines, Plates, and Datasets

We further evaluated drug molecular representation methods for predicting drug-induced gene expression under three generalization protocols: leave-cell-line-out, leave-plate-out, and leave-dataset-out (see **Methods: Evaluation Protocol**).

In the leave-cell-line-out evaluation on LINCS2020 (**Fig. 2e**), Chemprop achieved the highest DEG_PCC (median = 0.620), followed closely by Mole-BERT (median = 0.617) and InfoAlign (median = 0.615), with KPGT (median = 0.611) and UniMol (median = 0.612) performing slightly lower. Notably, the ECFP4 method remained highly competitive (median = 0.605), indicating that rule-based molecular representations still provide a robust benchmark for cross-cell line prediction. The corresponding DEG_RMSE metrics revealed a consistent ranking pattern with the DEG_PCC results (**Supplementary Fig. 2c**).

In the leave-plate-out evaluation on Tahoe_mini, we trained models on a single plate and evaluated on all remaining plates. Based on transfer difficulty (**Supplementary Note 1, Fig. 2f**), plates P1 and P13 were shown as challenging outlier sources (**Supplementary Fig. 2f**). When trained on the outlier P1 plate (**Fig. 2f**, left), all models failed to generalize effectively, with DEG_PCCs values close to random performance (Ouroboros/MolCLR: 0.109; ECFP4: 0.107). However, models trained on plate P13 demonstrated improved overall generalization performance (**Fig. 2f**, right). UniMolV2 achieved the highest performance (DEG_PCC = 0.405), but its advantage over ECFP4 (0.404) was negligible, with most methods falling within a narrow range (±0.01; **Supplementary Fig. 2d-e**). These results indicate that advanced molecular representations provide limited robustness to experimental batch effects, offering little advantage over simple fingerprint-based methods.

Cross-dataset transfer between LINCS2020 and Tahoe_mini represents the most challenging generalization scenario (**Fig. 2g**). To ensure fair comparisons, all models were evaluated on the 965 genes shared between the two datasets. Due to substantial inter-domain inconsistency (**Supplementary Note 1**), transfer from LINCS2020 to Tahoe_mini resulted in a near-complete performance collapse, with the best-performing ECFP4 reaching a DEG_PCC of only 0.01 (**Fig. 2g**, left). Reverse transfer from Tahoe_mini to LINCS2020 showed modest improvement but remained far below in-domain benchmarks, with GeminiMol and Mole-BERT achieving approximately 0.11 (**Fig. 2g**, right). Collectively, these results indicate that current representation methods degrade severely under stringent generalization conditions. In contrast, the canonical ECFP4 fingerprint remains comparatively robust, often matching or outperforming more complex deep learning-based molecular representations.

### Benchmarking Drug Molecular Representations for Predicting Drug-Induced Morphology Phenotypes

Cell morphology serves as a critical phenotypic readout, capturing the integrated structural and functional responses of cells to drug treatments^33,54,55^. We next conducted a systematic benchmark of 12 drug molecular representation methods for predicting drug-induced cellular morphology (**Fig. 1d**, *STP-Drug representations focus*). This analysis evaluates how different molecular encodings of drug compound affect the prediction of post-treatment morphological phenotypes across three high-content imaging datasets (cpg0016, BBBC036_Image and BBBC047_Image) under a leave-SMILES-out setting that simulates prediction for unseen compounds.

On the cpg0016 dataset, performance differences among representation methods were subtle but discernible (**Fig. 3a**). Chemprop achieved the highest correlation (median DEG_PCC = 0.344), followed closely by UniMolV2 (0.341) and Mole-BERT (0.339). Notably, Chemprop outperformed the standard ECFP4 baseline (median DEG_PCC = 0.318) by approximately 8.3% and also achieved the lowest error (median DEG_RMSE = 0.837). Direction_ACC results further confirmed this advantage, with Chemprop (0.607) leading UniMolV2 (0.606) and Mole-BERT (0.605).

**Fig. 3:**
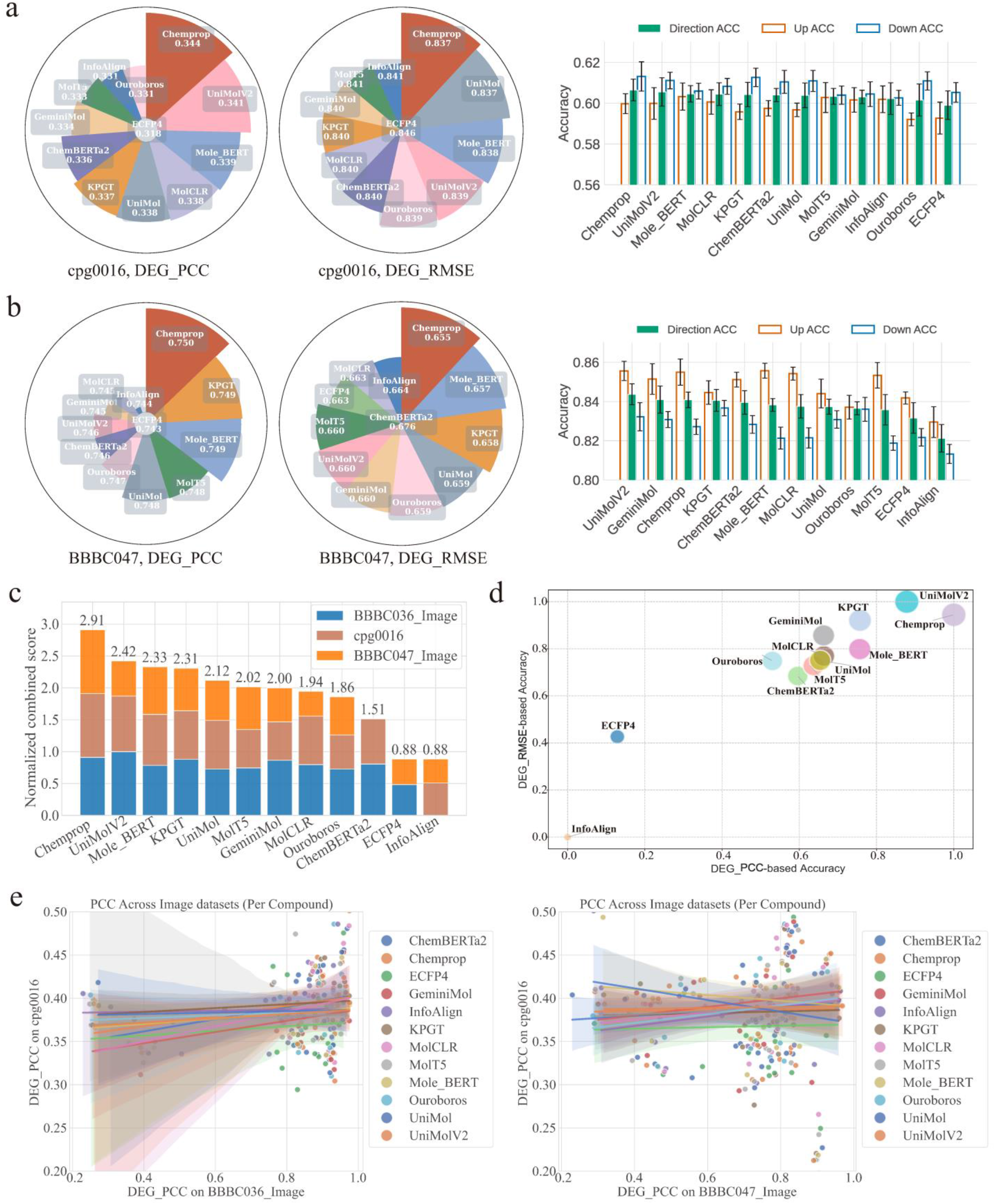
Benchmark drug molecular representation methods in predicting drug-induced cell morphology. **a**, Results on the cpg0016 dataset. DEG_PCC, DEG_RMSE and Direction_ACC were used to evaluate prediction accuracy and directional consistency. **b**, Results on the BBBC047_Image dataset. **c**, Unified ranking of 12 molecular representation methods across three diverse imaging datasets (cpg0016, BBBC036_Image, and BBBC047_Image). **d**, Aggregate cross-dataset performance represented as a bubble chart integrating DEG_PCC and DEG_RMSE. **e**, Cross-dataset consistency of compound-wise prediction performance. Scatter plots show compound-level DEG_PCC scores compared across imaging datasets, including cpg0016 versus BBBC036_Image (left) and cpg0016 versus BBBC047_Image (right).

On the BBBC047_Image dataset, performance differences between methods were relatively small (**Fig. 3b**). Although Chemprop achieved the highest median DEG_PCC (0.750), its performance was comparable to KPGT, Mole-BERT, MolT5, and UniMol. In this setting, deep learning-based methods showed a modest but consistent advantage over the ECFP4 baseline (0.743), with improvements of approximately 2–3%, indicating their enhanced ability to capture morphological phenotypes. On the BBBC036_Image dataset, UniMolV2 achieved the best overall performance, with Chemprop ranking among the top five. Both methods substantially outperformed the ECFP4 baseline (**Supplementary Fig. 3**).

To evaluate performance variability across datasets (cpg0016, BBBC047_Image, and BBBC036_Image), we performed a comparative analysis by integrating normalized DEG_PCC and DEG_RMSE metrics (**Fig. 3c**). To ensure fair comparison, all metric values were min-max normalized and aggregated into a combined score. This analysis revealed a clear separation in performance among the evaluated methods. Chemprop demonstrated the most stable performance across all three datasets (score=2.91), followed by UniMolV2 (2.42), suggesting that learned molecular representations capture more stable biological signals across datasets. In contrast, the fingerprint-based baseline ECFP4 performed substantially worse (0.88). In addition, **Fig. 3d** summarizes model performance using normalized DEG_PCC and normalized DEG_RMSE. UniMolV2 (normalized DEG_PCC = 0.88, normalized DEG_RMSE = 1.00) and Chemprop (1.00, 0.94) lie near the top-right region, indicating high predictive accuracy and low error. Conventional methods such as ECFP4 (0.13, 0.42) and InfoAlign fall in the bottom-left region, reflecting weaker performance and higher error. Although the overall differences are moderate, advanced molecular representation methods consistently outperform baseline methods in morphology prediction.

Next, we examined whether model performance is consistent across different imaging platforms. **Fig. 3e** shows drug compound-wise comparisons of prediction performance (DEG_PCC) across imaging datasets. Each dot represents a shared compound, and colors indicate different drug molecular representation methods. The left panel compares cpg0016 with BBBC036_Image, while the right panel compares cpg0016 with BBBC047_Image. In both comparisons, the points are widely scattered and the trend lines are nearly flat, indicating little to no relationship between performance across datasets. In other words, a compound that is predicted well in one dataset is not necessarily predicted well in another. These results suggest that morphology prediction is strongly affected by platform-specific factors, and that models trained on a single imaging dataset have limited ability to generalize across different platforms.

### Benchmarking Gene Representations for Predicting Drug-Induced Gene Expression

We next conducted a systematic benchmark of 12 gene representation methods for predicting drug-induced gene expression while holding the drug input fixed (**Fig. 1d**, *STP-Gene representations focus*). This aims to assess how different gene representation methods influence post-treatment transcriptomic predictions. Specifically, we compared a simple Raw_Gene baseline, in which raw gene expression profiles are fed directly into the predictor without any intermediate encoding, with 11 pretrained advanced models including tGPT, scFoundation, scGPT, Geneformer, CellPLM, scBERT, OpenBioMed, UCE, STATE, Cell2Sentence, and SCimilarity. These models first transform gene expression profiles into gene representations, which are then provided as inputs to the same MVCBench predictor.

On the large-scale LINCS2020 dataset, tGPT and STATE achieved the highest performance **(Fig. 4a)**, both reaching a DEG_PCC of 0.614 (±0.121 and ±0.119, respectively). They outperformed the next-best method, scFoundation (0.604 ± 0.122), by approximately 1.6%. Notably, scGPT (0.386 ± 0.138) and Cell2Sentence (0.385 ± 0.134) did not surpass the Raw_Gene baseline (0.603 ± 0.124). Similar trends were observed for DEG_RMSE. We further evaluated these gene representation methods on a subset of the top-100 highly expressed genes (HEGs) **(Fig. 4b)**. STATE again showed top-tier performance, consistently outperforming the Raw_Gene baseline. The same conclusion was observed on the Tahoe_mini P1 dataset (**Supplementary Fig. 4a**).

**Fig. 4:**
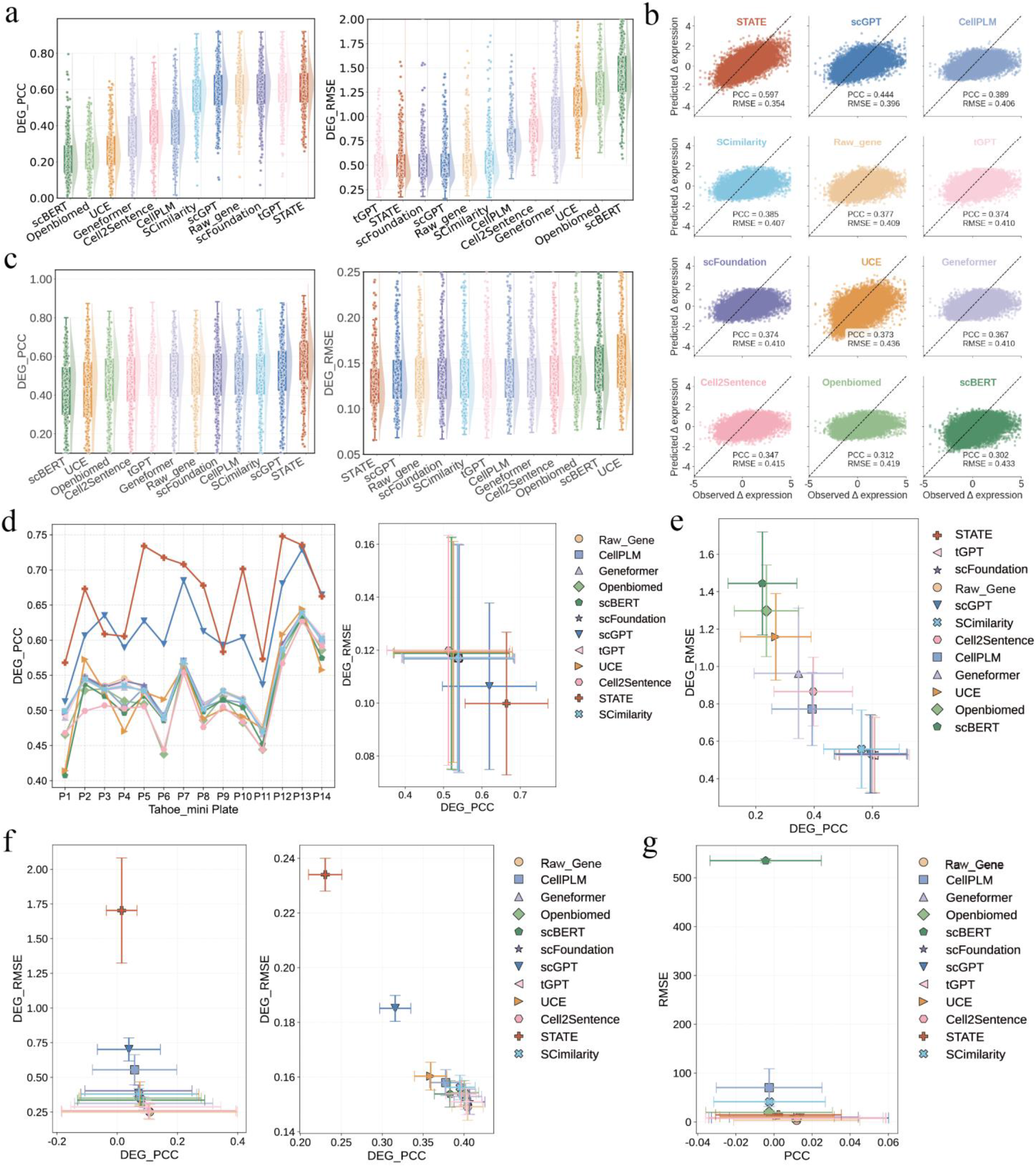
Benchmark gene representation methods for drug-induced gene expression prediction. **a**, Performance evaluation on LINCS2020 dataset. Comparison of 12 gene representation methods under the leave-SMILES-out strategy. **b**, Evaluation on the top-100 highly expressed genes (HEG) subset of LINCS2020. This subset comprises the top 100 genes with the highest mean basal expression in control samples, focusing on the reconstruction of high-signal biological features. Scatter plots illustrate the correlation between observed and predicted expression (drug-induced changes) specifically for these genes. c, Benchmarking on the Tahoe_mini P1 dataset. **d**, Robustness of gene representation models across all 14 Tahoe_mini plates. **e**, Leave-cell-line-out generalization on LINCS2020. **f**, Leave-plate-out generalization on Tahoe_mini. **g**, Leave-dataset-out generalization between LINCS2020 and Tahoe_mini.

We then evaluated gene representation methods on the recently released Tahoe_mini dataset. On plate P1, STATE achieved the best performance with a DEG_PCC of 0.579 ± 0.146 (**Fig. 4c**), while scGPT was the closest competitor with a DEG_PCC of 0.533 ± 0.164, showing clear improvement compared to its performance on LINCS2020. Evaluation across all 14 Tahoe_mini plates confirmed this pattern (**Fig. 4d**). STATE consistently outperformed other models and achieved a DEG_PCC of 0.664 ± 0.108, corresponding to a 22.7% improvement over the Raw_Gene baseline.

### Generalization of Gene Representations across Cell Lines, Plates, and Datasets

To evaluate the robustness of gene representation methods in predicting drug-induced gene expression, we evaluated them under three challenging generalization settings: (1) leave-cell-line-out; (2) leave-plate-out; and (3) leave-dataset-out (see **Methods: Evaluation Protocol**).

Under the leave-cell-line-out setting on LINCS2020 (**Fig. 4e**), STATE (DEG_PCC = 0.606 ± 0.119) and tGPT (DEG_PCC = 0.605 ± 0.121) achieved the highest correlations, leading a top tier that included scFoundation (DEG_PCC=0.597 ± 0.123), Raw_Gene (DEG_PCC=0.596 ± 0.122), and scGPT (DEG_PCC=0.593 ± 0.124). SCimilarity formed a distinct middle tier (DEG_PCC=0.562 ± 0.129). Other methods, including Cell2Sentence, CellPLM, Geneformer, UCE, OpenBioMed and scBERT, performed substantially worse. DEG_RMSE values showed a similar trend, with tGPT achieved the lowest error (0.524 ± 0.202), outperforming both Raw_Gene (0.534 ± 0.203) and scFoundation (0.529 ± 0.207).

Under the leave-plate-out setting on Tahoe_mini, we focused on plates P1 and P13, which represent challenging outlier scenarios due to strong batch effects and low similarity to other plates (**Supplementary Note 1** and **Supplementary Fig. 1c**). When generalizing from the distinct P1 plate, the simple Raw_Gene baseline proved surprisingly resilient (**Fig. 4f**, left; **Supplementary Fig. 4b**, top), achieving the highest correlation (DEG_PCC = 0.107 ± 0.291). It was followed closely by Cell2Sentence (0.104 ± 0.289) and tGPT (0.095 ± 0.249). In contrast, STATE, which otherwise excels in biological transfer, dropped to near the bottom (DEG_PCC = 0.014 ± 0.050), performing worse than scGPT (0.038 ± 0.104). DEG_RMSE rankings mirrored this trend, with Raw_Gene achieving the lowest error (DEG_RMSE = 0.251 ± 0.054). Although overall performance improved when models were trained on P13 (**Fig. 4f**, right; **Supplementary Fig. 4b**, bottom), foundation models still did not consistently outperform the baseline. This result suggests that more complex models are more susceptible to technical overfitting than simpler representations under strong batch effects.

Finally, in the leave-dataset-out setting, where models trained on LINCS2020 were evaluated on Tahoe_mini, predictive performance was severely reduced across all methods (**Fig. 4g**). Within this challenging scenario, scGPT achieved the highest correlation (DEG_PCC = 0.013 ± 0.046), although overall performance remained very low. When models trained on Tahoe_mini were evaluated on LINCS2020 (**Supplementary Fig. 4c**), tGPT and Geneformer achieved the best performance (DEG_PCC=0.129 ± 0.09), surpassing the Raw_Gene baseline (DEG_PCC=0.080± 0.07). Analysis of transfer difficulty confirmed substantial platform-specific shifts (Alter_PCC = 0, Alter_RMSE = 0.24, **Supplementary Fig. 1c**). A combined view integrating all metrics (**Supplementary Fig. 4d**) shows that STATE and scGPT consistently achieve relatively higher correlation and lower error, indicating that they are among the most effective gene representation methods for predicting drug-induced gene expression.

### Benchmarking Drug Molecular Representations for Joint Prediction of Drug-Induced Gene Expression and Morphology

To move beyond single-task paradigm (STP; i.e. predicting drug-induced gene expression or cellular morphology), we further conducted a systematic evaluation of drug molecular representation methods within the multimodal virtual cell (MVC) paradigm, for the joint prediction of drug-induced gene expression and cellular morphology. This analysis aims to assess how different molecular encodings of drug compounds influence the simultaneous prediction of post-treatment transcriptomic and morphological responses (**Fig. 1d**, *MVC* paradigm). Using the BBBC047 and BBBC036 datasets, models were trained to predict paired drug-induced gene expression (GE) and Cell Painting morphology (CP) phenotypes from compound structures and pre-treatment cellular profiles.

On the BBBC047 dataset (**Fig. 5a**), UniMolV2 achieved the strongest overall performance across both modalities, reaching the highest accuracy for morphology prediction (CP: DEG_PCC = 0.736, CP: DEG_RMSE = 0.766) and also delivered the best performance for gene expression prediction (GE: DEG_PCC = 0.702, GE: DEG_RMSE = 0.959). In contrast, Chemprop, which performed competitively in several STP tasks, showed weaker performance in this MVC setting, trailing UniMolV2 by 3.2% in correlation and showing 4.5% higher prediction error. On the BBBC036 dataset (**Fig. 5b**), performance became more modality-dependent. For CP prediction, KPGT achieved the highest correlation (CP: DEG_PCC = 0.799), followed closely by UniMol and Ouroboros. However, Chemprop ranked first for gene expression prediction (GE: DEG_PCC = 0.717), while several structure-based molecular foundation models performed better on morphology than on transcriptional responses. **Fig. 5c** summarizes performance across both datasets (BBBC047 and BBBC036). InfoAlign consistently ranked among the weakest methods, indicating a limited ability to capture the coupled transcriptional and morphological responses induced by drug perturbations. Among the evaluated methods, UniMolV2 demonstrated the most consistent performance, making it a strong default choice for chemically informed multimodal virtual cell modeling.

**Fig. 5:**
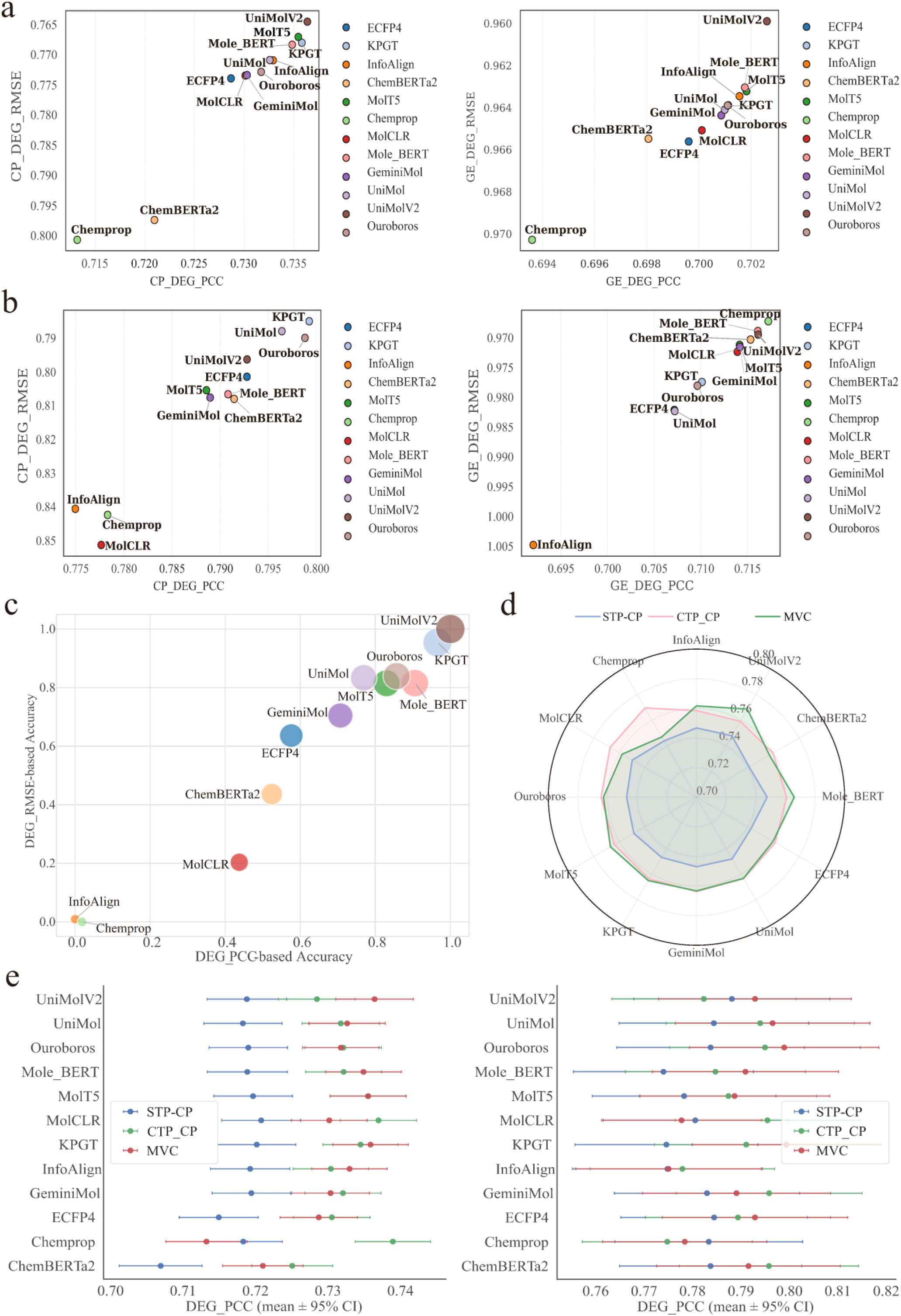
Benchmarking drug molecular representations for joint prediction of drug-induced gene expression and morphology. **a**-**b**, Performance of molecular representation methods on paired Cell Painting (CP) and gene expression (GE) prediction using the BBBC047 and BBBC036 datasets. **c**, Integrated ranking of molecular representations across datasets and modalities. **d**-**e**, Comparison of three modeling paradigms for CP prediction: single-target prediction (STP), cross-modal assisted prediction (CTP), and multimodal virtual cell (MVC) paradigm. STP uses only the pre-treatment profile of the target modality; CTP additionally incorporates the complementary pre-treatment modality as auxiliary input; MVC jointly predicts post-treatment CP and GE using a multitask objective. Corresponding GE results are shown in Supplementary Fig. 5a,b.

We next examined whether multimodal integration itself improves drug-induced phenotype prediction. To this end, we compared three modeling paradigms that differ in how multimodal information is incorporated. In the single-task paradigm (**Fig. 1d**, *STP* paradigm), the model predicts a single post-treatment modality using the drug representation and the corresponding pre-treatment profile of the target modality only (e.g. STP-CP: Drug + pre-CP → post-CP; STP-GE: Drug + pre-GE → post-GE). This setting isolates unimodal conditional prediction without cross-modal interactions. In the cross-modal assisted paradigm (*CTP*), the pre-treatment profile of the complementary modality is provided as additional context, but the model is still optimized to predict only one target modality (e.g. CTP-CP: Drug + pre-CP + pre-GE → post-CP; CTP-GE: Drug + pre-CP + pre-GE → post-GE). In the multimodal virtual cell (**Fig. 1d**, *MVC* paradigm, Drug + pre-CP + pre-GE → post-CP + post-GE) paradigm, both pre-treatment modalities are provided as input to a shared backbone, and the model jointly predicts post-treatment GE and CP through modality-specific output heads under a multitask objective.

Across molecular representations, MVC consistently outperformed STP for morphology prediction on both datasets (**Fig. 5d-e**), with relative performance gains of 2–6%. CTP also improved over STP in most cases, indicating that incorporating complementary pre-treatment information provides useful predictive context even without explicit joint optimization. A similar trend was observed for gene expression prediction (**Supplementary Fig. 5a**,**b**), suggesting that the benefits of multimodal integration are consistent across both phenotypic modalities. These results demonstrate that accurate multimodal virtual cell prediction depends not only on the quality of drug molecular representations but also on the effective incorporation of complementary pre-treatment modalities. Models that leverage multimodal context consistently outperform single-modality approaches, with the largest gains achieved under joint multimodal modeling.

### Design Principles for Multimodal Virtual Cell Architectures

Since **Fig. 5d-e** shows that incorporating multimodal information and joint supervision improves predictive fidelity relative to unimodal prediction, we therefore sought to understand why multimodal integration is beneficial and which architectural choices most strongly influence joint prediction of drug-induced transcriptomic and morphological phenotypes in the MVC paradigm.

As a first step, we investigated whether transcriptomic and morphological phenotypes provide redundant or complementary information about drug-induced cellular responses. If the two modalities encode overlapping biological signals, compounds predicted accurately in gene expression should also be predicted accurately in morphology. Herein, we quantified compound-wise rank correlations of DEG_PCC scores between gene expression prediction (LINCS 2020) and morphology prediction (cpg0016). These correlations were close to zero (**Fig. 6a**), indicating that compounds predicted accurately in the transcriptomic domain were not necessarily predicted well in the morphological domain, and vice versa. We further examined this relationship within the BBBC047 dataset (**Fig. 6b**), where similarly low correlations were observed. This suggests that predictive performance in gene expression provides little information about performance in morphology. A comparable pattern was also observed in the independent BBBC036 dataset (**Supplementary Fig. 6a**). To further validate this independence, we projected molecular, GE, and CP embeddings into a unified t-SNE space. Perturbations induced by the same compounds did not form coherent clusters across modalities (**Supplementary Fig. 6b**), indicating that these representations reside in distinct information subspaces. Together, these results demonstrate that transcriptomic and morphological readouts capture distinct and complementary, rather than redundant, aspects of drug-induced cellular responses.

**Fig. 6:**
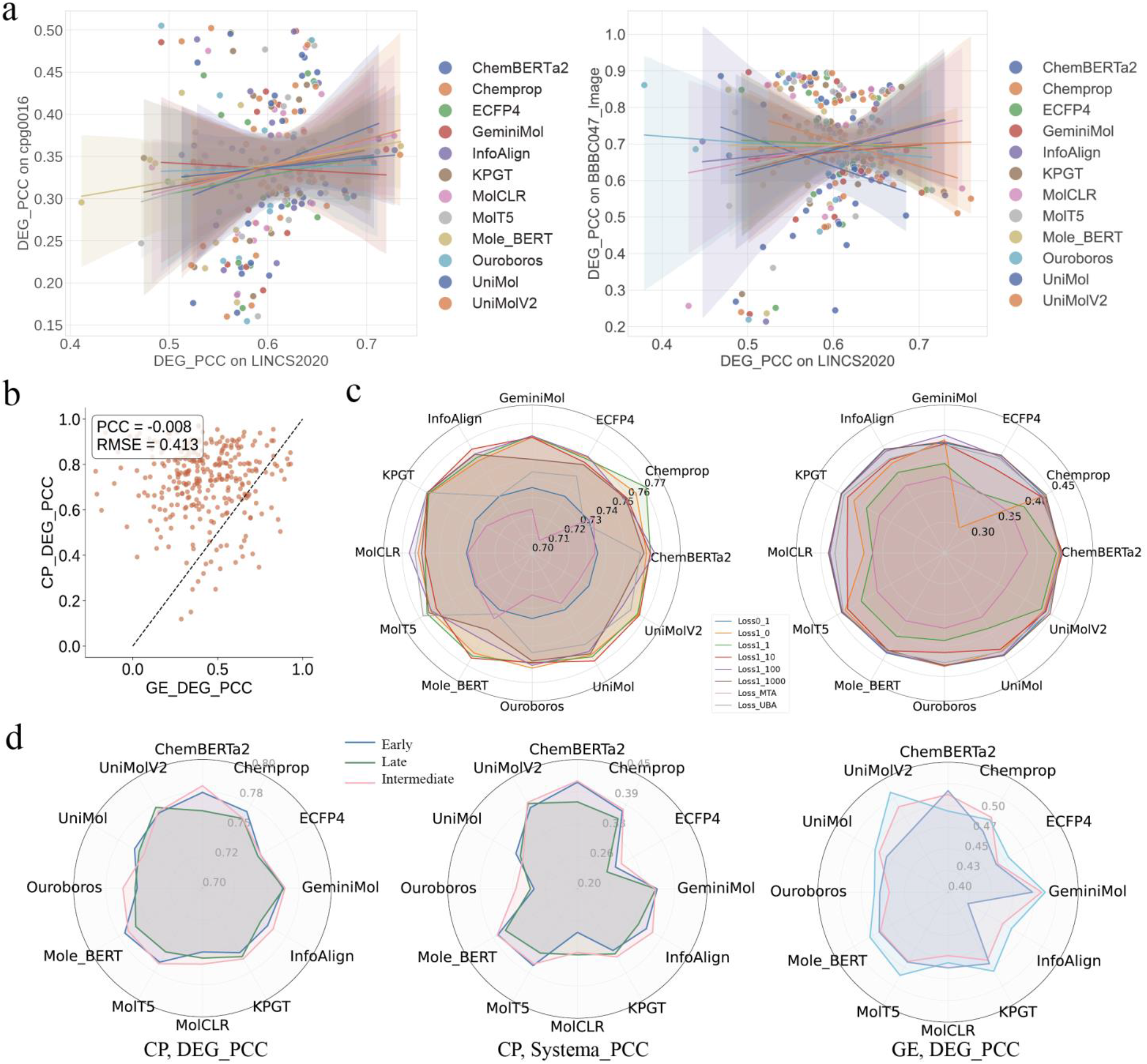
Optimization of multimodal virtual cell architectures. **a**, Compound-level correlations between gene expression and morphology prediction accuracy across datasets, assessing whether predictive performance transfers across modalities. **b**, Cross-modality performance landscape for individual compounds in BBBC047. Each point represents a compound positioned by its gene expression prediction accuracy (x axis) and morphology prediction accuracy (y axis). The near-zero correlation (PCC = −0.008) indicates minimal alignment of predictive performance across modalities. **c**, Evaluation of loss-weighting strategies on BBBC047. Left, morphology prediction peaks at an approximately balanced fixed-ratio loss (DEG_PCC = 0.733). Right, gene expression prediction reaches optimal performance when the transcriptomic loss is strongly up-weighted (DEG_PCC = 0.429). **d**, Comparison of early, intermediate and late fusion strategies for integrating transcriptomic and morphological information on BBBC047.

We next examined how these two modalities should be integrated during joint optimization in the MVC paradigm. Transcriptomic and morphological data differ substantially in measurement characteristics: gene expression profiles are sparse and prone to dropout, whereas morphological features are continuous and often influenced by batch-specific technical variation. As a result, the corresponding loss terms differ in both scale and gradient magnitude, which can bias the optimization of joint prediction. Therefore, we systematically compared fixed-ratio weighting schemes with adaptive loss strategies, including Multi-Task Adaptive (MTA) loss and Uncertainty-Based Adaptive (UBA) weighting loss (**Fig. 6c**; **Methods: Loss Function for MVC Architectures**). Fixed-ratio loss consistently outperformed adaptive loss on BBBC047. Morphology prediction was maximized at an approximately balanced 1:1 loss ratio (**Fig. 6c**, left; DEG_PCC = 0.733), whereas optimal gene expression prediction required substantial up-weighting of the transcriptomic loss, reaching its best performance when the GE loss was amplified 100-fold (**Fig. 6c**, right; DEG_PCC = 0.429). By contrast, both adaptive losses underperformed across settings, suggesting that their uncertainty-driven formulations do not adequately account for these two modalities.

We then evaluated three fusion strategies for integrating transcriptomic and morphological information in the MVC paradigm: early fusion, intermediate fusion, and late fusion (**Fig. 6d, Methods: Fusion Strategies**). Early fusion, which concatenates both modalities at the input level, performed worst overall. This result is expected due to the substantial heterogeneity between modalities, which likely limits the effectiveness of direct feature-level concatenation. Intermediate fusion, where the two modalities are integrated after unimodal encoding, achieved the strongest performance for morphology prediction (DEG_PCC = 0.73 ± 0.01; Systema_PCC = 0.32 ± 0.03). This suggests that combining modalities after extracting stable unimodal representations allows the model to better utilize complementary information. In contrast, late fusion, which integrates modality-specific predictions at the decision level, yielded the best performance for gene expression prediction (DEG_PCC = 0.439 ± 0.01). This pattern indicates that decision-level integration is more effective when transcriptomic prediction already benefits from strong unimodal predictors. Together, these results show that the optimal fusion depth depends on the prediction target rather than being universal across modalities.

## DISCUSSION

This study introduces MVCBench, a systematic framework for evaluating how drug molecular and gene representations influence the prediction of drug-induced cellular phenotypes. By integrating large-scale multimodal datasets capturing pre- and post-treatment states with a controlled benchmarking protocol, MVCBench enables comprehensive and fair comparisons of representation methods across both single-task (STP) and multimodal virtual cell (MVC) paradigms.

In the STP paradigm, our benchmarking yield several key insights into the current landscape of representation learning for drug perturbation modeling. First, the effectiveness of learned representations is highly dependent on the phenotypic modality being predicted. Advanced drug molecular representation methods, particularly geometry-aware models such as UniMolV2 and KPGT, consistently improve the prediction of drug-induced morphological phenotypes, yet offer only limited gains over classical fingerprints for gene expression prediction. Second, gene representation methods play a dominant role in transcriptomic prediction. Task-specific models, most notably STATE, substantially outperform general-purpose single-cell foundation models such as scGPT and scFoundation. This suggests that models trained specifically to capture post-treatment gene expression provide more informative embeddings than those optimized for general transcriptomic representation learning. Third, across both drug molecular and gene representation methods, predictive performance deteriorates markedly under distribution shift, including cross-cell-line, cross-batch, and cross-dataset settings. These findings highlight a key limitation for real-world deployment, as current models rely heavily on dataset-specific signals and struggle to generalize robustly across experimental platforms and biological contexts.

In the MVC paradigm, our analyses further demonstrate that incorporating both transcriptomic and morphological context consistently improved predictive performance relative to single-modality models (STP). These two modalities capture complementary aspects of drug-induced cellular responses rather than redundant signals. This complementarity likely reflects the multilayer nature of drug perturbation biology, in which transcriptional regulation and structural cellular remodeling arise through partially distinct regulatory pathways. Beyond benchmarking, we also provide practical design principles for multimodal virtual cell architectures. First, gene expression and cell morphology should be treated as complementary, non-redundant readouts of drug response. Second, effective multimodal optimization requires explicit calibration of modality-specific loss scales, rather than relying solely on adaptive weighting mechanisms. Third, fusion strategies should be selected according to the prediction target, with intermediate fusion favoring morphology prediction and late fusion favoring gene expression prediction.

Despite the comprehensive design of MVCBench, several limitations should be acknowledged. First, the availability of truly paired multimodal datasets remains limited. Most transcriptomic and morphological measurements are derived from separate assays, rather than collected from the same cells under identical conditions. This may confound multimodal integration and evaluation^56,57^. Second, although MVCBench systematically controls for representation inputs, it relies on a fixed predictive architecture. While this design enables fair comparison, it may not fully capture interactions between representation methods and model architectures, potentially underestimating the performance of certain methods that benefit from specialized model designs. Third, the current framework primarily focuses on transcriptomic and morphological modalities, without incorporating additional biological layers such as proteomics or metabolomics. Finally, the observed performance degradation under distribution shift highlights a broader limitation of current representation learning approaches. Although MVCBench exposes these generalization challenges, it does not yet provide solutions for improving robustness across datasets, experimental platforms, or biological contexts.

Looking forward, the development of standardized multimodal datasets, improved cross-domain representation learning methods and mechanistically informed modeling frameworks will be essential for advancing virtual cell prediction. By providing a systematic benchmarking foundation and revealing key design principles for MVC architectures, MVCBench represents a step toward building more robust and biologically meaningful models of drug-induced cellular responses.

## METHODS

### Drug Molecular Representation Methods

We evaluated 12 representative molecular representation methods spanning four major paradigms to cover the mainstream strategies in drug encoding. As a classical baseline, we included the rule-based fingerprint ECFP4^19^. For graph-based learning, we selected Chemprop^22^, MolCLR^34^, and InfoAlign^35^, representing message-passing architectures and contrastive pre-training on molecular graphs. We further examined Transformer-based models, including ChemBERTa2^36^, Mole-BERT^37^, MolT5^38^, and KPGT^23^, to assess sequence-driven and knowledge-guided pre-training approaches. Finally, to evaluate the contribution of explicit structural information, we benchmarked geometry-aware pre-trained models that incorporate conformational cues or structural constraints during representation learning, including UniMol^39^, UniMolV2^40^, GeminiMol^41^, and Ouroboros^42^. Detailed architectural configurations, pre-training objectives, and checkpoint information are provided in **Supplementary Note 2**.

### Gene Representation Methods

We benchmarked 12 gene representation strategies, including a raw gene baseline and 11 single-cell foundation models (scFMs). The scFMs were grouped according to their architectural backbones: 1) Encoder-only models: Geneformer^43^, scBERT^44^, OpenBioMed^45^, SCimilarity^46^, which use BERT-like objectives to learn contextual gene embeddings. 2) Encoder-decoder models: scFoundation^47^, scGPT^18^, UCE^48^, CellPLM^49^, STATE^50^, designed for generative tasks and cross-modal translation. 3) Decoder-only models: tGPT^51^ and Cell2Sentence^52^, which follow autoregressive generative paradigms. All models were evaluated with frozen weights to assess the practical utility of their learned representations under a shared lightweight predictor. Full implementation details are provided in **Supplementary Note 2**.

To ensure strict comparability across architectures, static embeddings from all single-cell foundation models (with frozen weights) were fed into an identical MLP for perturbation prediction. This setup decouples representation learning from task-specific modeling, isolating the contribution of the embeddings. However, reported performance reflects the combined effect of the foundation model and the shared downstream predictor. Because the MLP is trained on the evaluation dataset, residual batch effects or dataset-specific biases may influence results, especially in cross-domain scenarios. Therefore, these benchmarks should be interpreted as evaluating the practical utility of model embeddings when used with a lightweight predictor, rather than as a pure measure of intrinsic representation quality.

### MVCBench Framework

As illustrated in **Fig. 1d**, the MVCBench framework is organized around two main research pipelines: the single-task paradigm (STP) with drug/gene representation focus pipeline, which predicts either gene expression or cell morphology independently, and the multimodal virtual cell paradigm, implemented via the MVC architecture for joint prediction of both phenotypes. In both pipelines, we adopt a minimalist architecture to reduce confounding factors and ensure rigorous benchmarking. Implementation details for each pipeline are described below.

#### 1) Single-task paradigm (STP): Drug/Gene representation focus

To standardize evaluation across the independent component benchmarking tasks, specifically the STP-Drug Representations Focus and STP-Gene Representations Focus outlined in **Fig. 1d**, we implemented a lightweight single-task perturbation predictor. This controlled evaluation head ensures that performance differences reflect the quality of the input representations rather than downstream model complexity. The predictor follows a conditional encoder-decoder design, predicting a single post-treatment modality (either gene expression or cell morphology) by integrating variable molecular features with the fixed pre-treatment cellular state.

Formally, let ***x***^(0)^ ∈ ℝ^*d*^ denote the baseline cell profile, representing either transcriptomic or morphological features, and ***h***_*d*_ ∈ ℝ^*k*^ denote the drug molecular embedding. The encoder maps the baseline profile into a latent representation ***z***_*cell*_ using a single fully connected layer with batch normalization and a non-linear activation:

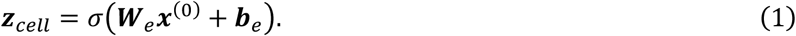

To incorporate drug information, the drug embedding ***h***_*d*_ is concatenated with ***z***_*cell*_ to form a joint representation z_*joint*_:

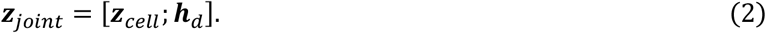

The decoder, comprising a linear layer with activation, maps ***z***_*joint*_ to the predicted perturbed state 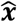:

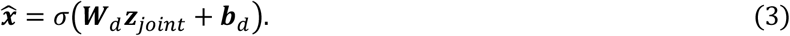

For single-task benchmarking, the model is trained using mean squared error (MSE) between 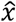 and the observed perturbed profile *x*_gt_, providing a consistent basis for comparing representation quality.

#### 2) Multimodal virtual cell (MVC) paradigm

To implement the MVC outlined in **Fig. 1d**, we designed a dual-encoder, dual-decoder network. This architecture benchmarks drug molecular representations while optimizing joint perturbation prediction strategies, including fusion depth and loss weighting (**Fig. 6**). The network consists of six fully connected layers organized into three functional modules: two modality-specific encoders (2 layers) for independent feature extraction, two fusion layers (2 layers) for cross-modal integration, and two task-specific decoders (2 layers) for predicting the respective phenotypes.

To distinguish between modalities, let 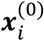and 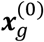 denote the baseline cell morphology and gene expression profiles respectively, while ***h***_*d*_remains the drug molecular vector. The morphology encoder *E*_*i*_ and expression encoder *E*_*g*_ first map their respective inputs into latent representations:

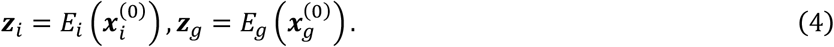

These representations are concatenated with ***h***_*d*_ to form a fused vector:

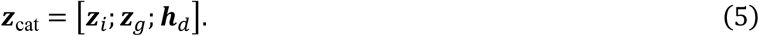

Two additional MLP layers refine ***z***_cat_ into a unified representation ***h***_fused_:

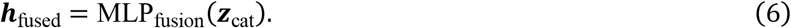

Finally, ***h***_fused_ is passed to two task-specific decoders to predict post-treatment profiles:

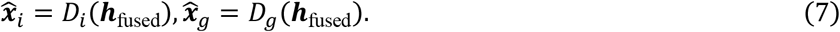

All fully connected layers (except input/output) include batch normalization and non-linear activation, consistent with the baseline configuration. The multitask objective minimizes the weighted sum of reconstruction errors:

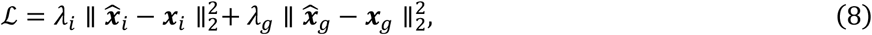

where ***x***_*i*_ and ***x***_*g*_ are the observed post-treatment profiles, and *λ*_*i*_, *λ*_*g*_ are loss weights. Unless otherwise specified, the multimodal model used a fixed loss ratio of 1:100 for CP and GE, selected based on validation experiments described in the Results section. This design aligns baseline cellular states and molecular features in a shared latent space through independent encoding, intermediate fusion, and multitask decoding, enabling effective joint prediction of morphological and transcriptomic responses without excessive architectural complexity.

#### 3) Loss function for MVC architectures

While MSE serves as the foundational loss for both single-task and multimodal prediction, the MVC framework requires additional strategies to balance learning signals across modalities with different scales and noise levels. We systematically evaluated three weighting schemes: Fixed-ratio weighting; Multi-task adaptive loss^58^; Uncertainty-weighted multimodal loss. For fixed-ratio weighting, we applied static scalar weights (e.g., *λ*=1, 10, 100) to manually calibrate the contribution of each modality. For multi-task adaptive loss, we implemented a dynamic weighting scheme to address the imbalance where one task dominates gradient updates. Let ℒ_*g*_ and ℒ_*i*_ denote the individual MSE losses for the gene expression and cell morphology prediction tasks respectively. The task weights are computed inversely proportional to the loss of the competing modality:

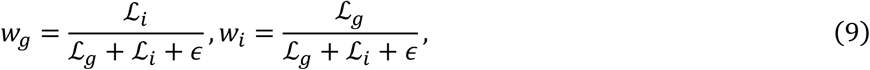

where *ϵ* is a small constant to prevent division by zero. The resulting adaptive multi-task loss is

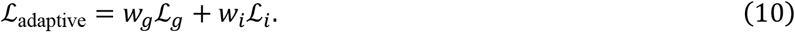

This approach effectively downweights tasks with higher losses (presumably harder tasks or those with larger scales), preventing them from dominating model updates and encouraging balanced learning across modalities.

For uncertainty-weighted multimodal loss, we implemented an uncertainty-weighted loss inspired by homoscedastic uncertainty estimation, to jointly optimize gene expression and cell morphology prediction. This formulation treats task-dependent uncertainty as a learnable scalar. Two parameters, *σ*_*g*_ and *σ*_*i*_, encode the relative uncertainty (noise variance) of the gene and morphology tasks respectively. The combined loss is defined as:

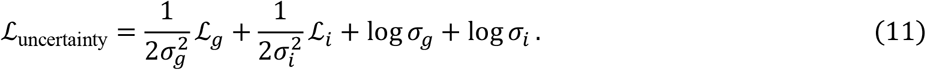

Intuitively, tasks with higher predictive uncertainty (larger *σ*) contribute less to the total loss, allowing the model to adaptively balance learning between modalities. During training, both *σ*_*g*_ and *σ*_*i*_ are learned jointly with model parameters, enabling automatic task weighting without manual hyperparameter tuning.

### Benchmarking Datasets and Preprocessing

We employed six benchmark datasets spanning transcriptomic, morphological, and multimodal profiles (detailed in **Table 1**). To ensure clarity, we adopt the following nomenclature for the multimodal datasets (CDRP-BBBC047-Bray and CDRPBIO-BBBC036-Bray). 1) BBBC047 and BBBC036 refer to the paired multimodal datasets used for joint training. 2) The *suffix* _Gene (e.g., BBBC047_Gene) denotes the transcriptomic component only. (3) The *suffix* _Image (e.g., BBBC047_Image) denotes the morphological component only.

#### Gene expression datasets

We curated five datasets including LINCS2020^13,30^, Tahoe-100M^31^ (comprising 14 subsets), CIGS^32^, BBBC047_Gene, and BBBC036_Gene. Collectively, these resources encompass 38,950 compounds and 482,323 paired pre- and post-treatment transcriptomic profiles. LINCS2020, BBBC047_Gene BBBC036_Gene were generated using the L1000 profiling platform, with raw measurements covering 978 landmark genes. Tahoe-100M dataset leverages the Mosaic high-throughput platform in combination with Parse Biosciences’ GigaLab for single-cell sample preparation, covering approximately 20,000 genes. To enable rigorous cross-dataset evaluation, we constructed Tahoe_mini by filtering the original gene space to the 965 landmark genes shared with LINCS2020, providing a dimensionally aligned basis for domain adaptation experiments. The CIGS dataset employs HTS and HiMAP-seq technologies to quantify 3,407 key genes. Notably, CIGS was recently released and was not included in the pretraining of any representation models, ensuring unbiased evaluation of out-of-domain generalization.

To ensure consistency across datasets, we implemented a unified preprocessing pipeline. Taking the CIGS dataset as an example, raw expression matrices were first separated into metadata and numeric components to restrict subsequent operations to expression values. Expression counts were then normalized by library size and scaled to counts per million (CPM-like) to mitigate sequencing depth variability, using the formula: 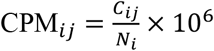, where *C*_*ij*_ denotes the raw count of gene *j* in sample *i*, and *N*_*i*_ is the total library size for sample *i*. Normalized values were log-transformed as log_2_(*x* + 1) to stabilize variance and compress dynamic range, reducing the influence of extreme values. Samples with zero or near-zero total counts were removed prior to normalization to ensure data quality.

#### Cell morphology datasets

We leveraged high-dimensional morphological profiles derived from the Cell Painting assay, encompassing cpg0016^33^ (currently the largest public resource) and the morphological components of the BBBC datasets (BBBC047_Image, BBBC036_Image). Collectively, these resources cover 84,858 compounds and provide 601,412 paired pre- and post-treatment profiles.

To prepare these data for downstream modeling, we implemented a standardized processing pipeline. First, raw single-cell features were aggregated at the well level by computing the mean across all cells, yielding replicate-level profiles. Next, to mitigate batch effects and ensure cross-plate comparability, we standardized these profiles within each plate (zero mean, unit variance) before averaging across replicate wells to obtain robust treatment-level profiles. This process resulted in final feature dimensions of 737 for cpg0016, 591 for BBBC036_Image, and 745 for BBBC047_Image.

#### Multimodal cell phenotypic datasets

BBBC047 and BBBC036 datasets were included as multimodal resources, containing 20,341 and 1,917 compounds, respectively. Given the substantially larger compound coverage in BBBC047, we use this dataset as an illustrative example. The BBBC047 dataset provides both transcriptomic and morphological profiles under compound perturbations. Following the preprocessing pipelines described above, we constructed matched five-element tuples of the form (compound SMILES, pre-treatment gene expression, pre-treatment morphology, post-treatment gene expression, post-treatment morphology) by averaging replicate measurements for each compound. These tuples formed the basis for multimodal fusion experiments. Prior to modeling, all modality-specific data were normalized, ensuring balanced contributions from both transcriptomic and morphological features.

### Evaluation Metrics

Accurate evaluation of perturbation response prediction requires metrics that simultaneously characterize structural agreement, magnitude fidelity, robustness to systematic variation, and biological interpretability. MVCBench therefore adopts a five-metric evaluation framework: DEG_PCC, DEG_RMSE, Systema_PCC^53^, Systema_RMSE, and Direction_ACC.

We first evaluate reconstruction quality using two standard differential-signature metrics. DEG_PCC measures the Pearson correlation between predicted and ground-truth perturbation-induced signatures, quantifying structural similarity of response profiles independent of absolute scaling. Complementarily, DEG_RMSE computes the root mean squared deviation between predicted and true signatures, capturing estimation bias in response magnitude. Together, these metrics assess whether a model recovers both the shape and intensity of perturbation effects relative to matched controls.

Formally, for each perturbation *p* defined over a feature space of size *G* (genes or morphological descriptors), let *x*_*p*_ denote the centroid of perturbed cells, *x*_*c*_ the matched control centroid, and 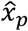 the predicted perturbed profile. The ground-truth and predicted differential signatures are defined as

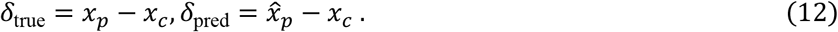

For the *i*-th perturbation, the corresponding signatures are denoted as 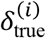 and 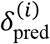. DEG_PCC and DEG_RMSE are computed over these differential signatures across all *N* perturbations.

However, metrics anchored to matched controls can be inflated by systematic domain shifts such as batch or plate effects. These shifts introduce global offsets shared across perturbations within a dataset or cell line, potentially leading to overoptimistic in-domain evaluation and degraded cross-domain generalization.

#### Systema_PCC and Systema_RMSE

To mitigate this bias, we adopt a centroid-referenced evaluation strategy inspired by the Systema^53^ framework. Instead of referencing matched controls, we define a domain-level centroid:

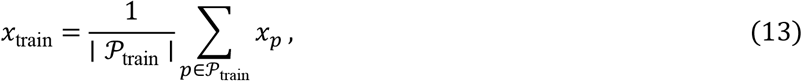

where *P*_train_ denotes all perturbations in the training set within the same domain. Thus, *x*_train_ captures the intrinsic geometric center of perturbation-induced states in that domain. Bias-corrected signatures are then defined as

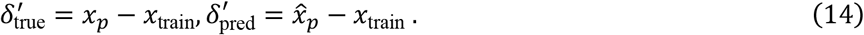

Under this formulation, Systema_PCC evaluates structural agreement in the bias-corrected perturbation space:

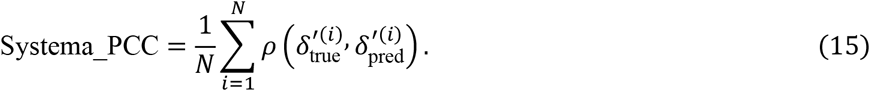

Similarly, Systema_RMSE extends DEG_RMSE to the bias-corrected setting:

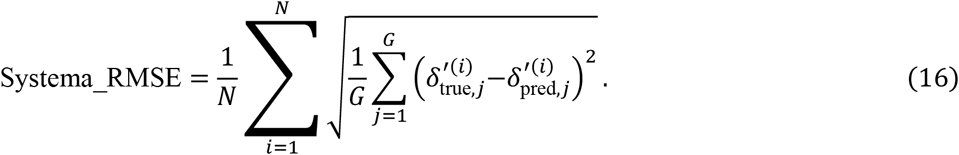

While Systema_PCC evaluates structural consistency after removing global domain offsets, Systema_RMSE measures magnitude fidelity of perturbation-specific responses under the same correction.

#### Direction_ACC

This metric provides an interpretable measure of sign consistency in feature regulation. For each perturbation *i*, we define a subset

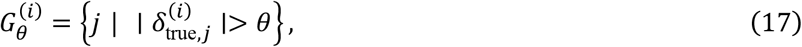

where *θ* > 0 excludes negligible changes. Directional accuracy is defined as:

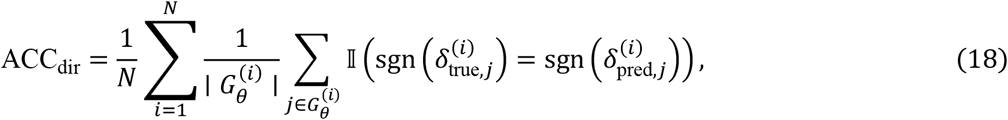

where 𝕀(·)denotes the indicator function. This metric captures agreement in regulatory direction independent of effect size, offering a biologically interpretable complement to correlation- and error-based measures.

### Evaluation Protocol

To rigorously assess model robustness and simulate real-world drug discovery scenarios, we implemented a hierarchical generalization protocol comprising three distinct levels of increasing difficulty: 1) Leave-SMILES-out: This serves as our primary evaluation baseline. Samples were stratified based on unique molecular structures (SMILES). Compounds present in the test set were strictly excluded from the training and validation sets, forcing the model to generalize to unseen chemical spaces. 2) Leave-plate-out and leave-cell-line-out: To evaluate robustness against biological variability, we held out specific experimental batches (plates) or cell lines during training, testing the model’s ability to transfer predictions to unseen biological environments. 3) Leave-dataset-out: The model was evaluated on completely independent datasets (e.g., external public datasets) not seen during model training, representing the highest level of transferability.

### Training Strategy

To ensure robust performance, we optimized the training configuration for the MVCBench MLP-based model. All models were trained using the Adam^59^ optimizer to minimize the mean squared error (MSE) between predicted and observed post-treatment profiles. The learning rate was set to 1×10^−3^ with a weight decay of 1×10^−5^ to regularize large weights and mitigate overfitting. Training was conducted for 1,000 epochs with a batch size of 1,000, and the random seed was fixed at 3407 to ensure reproducibility. The architecture consistently employed a two-layer encoder–decoder design, with hidden dimensions adjusted by task: 1,024 for single-modality prediction (512 per modality) and 1,536 for multimodal joint prediction. Additionally, inference latency benchmarks (Table 2) were conducted on a workstation equipped with an AMD Ryzen 9 7900X CPU and an NVIDIA GeForce RTX 4090 GPU (24GB VRAM) with a batch size of 32.

To ensure a rigorous comparison of the intrinsic representational quality of different representation models, the weights of all pre-trained molecular and gene models were kept strictly frozen during the training of the downstream predictor. The gradient updates were applied exclusively to the parameters of the specific predictor heads (i.e., the MLP encoders and decoders described in the Architecture sections), treating the foundation model outputs as static feature extractions. The best model checkpoint was selected based on validation-set DEG_PCC objective.

### Fusion Strategies

To rigorously benchmark the impact of information integration stages, we evaluated three fusion strategies: early, intermediate, and late fusion. Crucially, to ensure fair comparison, we maintained a consistent network depth of six fully connected layers (excluding input/output projections) across all architectures, varying only the fusion point and the distribution of shared versus modality-specific layers. 1) Intermediate fusion (default): As described above, this architecture employs a balanced design comprising 2 modality-specific encoder layers, followed by concatenation, 2 shared fusion layers, and 2 task-specific decoder layers (2+2+2 configuration). This allows for partial feature extraction prior to cross-modal interaction. 2) Early fusion: Input features from drug, gene, and morphology modalities are concatenated immediately at the input level. The unified vector is processed by 4 shared encoding layers to capture low-level cross-modal correlations, followed by 2 task-specific decoder layers for prediction (4+2 configuration). 3) Late fusion: Modalities are processed independently through deeper 4-layer modality-specific encoders to extract high-level semantic features. These high-level representations are concatenated and passed through 2 shared decoder layers to generate the final predictions (4+2 configuration). This design ensures that the longest path from input to output remains constant at six layers for all variants, attributing performance differences solely to the timing of information fusion rather than model complexity.

## DATA AVAILABILITY

All datasets used in this study are publicly available from established community resources. Transcriptomic profiles were obtained from LINCS2020 (L1000 platform), Tahoe-100M, CIGS, and the gene expression subsets of CDRP-BBBC047-Bray and CDRPBIO-BBBC036-Bray, collectively covering 38,950 compounds and 482,323 paired pre- and post-treatment transcriptomic profiles across multiple cell lines. Morphological features were derived from Cell Painting datasets, including cpg0016 and the morphology subsets of CDRP-BBBC047-Bray and CDRPBIO-BBBC036-Bray, encompassing 84,858 compounds and 601,412 paired morphological profiles. Multimodal resources include BBBC047 and BBBC036, which provide matched transcriptomic and morphological readouts under compound perturbations. All raw and processed data, along with metadata and preprocessing scripts, are accessible through their respective repositories: (1) LINCS2020: https://clue.io/data/CMap2020; (2) BBBC datasets: https://bbbc.broadinstitute.org; (3) cpg0016: https://github.com/broadinstitute/cellpainting-gallery/blob/main/README.md; (4) Tahoe-100M: https://huggingface.co/datasets/tahoebio/Tahoe-100M; (5) CIGS: https://cigs.iomicscloud.com. Detailed preprocessing pipelines and benchmark splits are provided in the MVCBench GitHub repository: https://github.com/QSong-github/MVCBench.

## CODE AVAILABILITY

A comprehensive project website showcasing key findings and benchmarks is available at https://QSong-github.github.io/MVCBench. All source code for data preprocessing, model implementation, training, and evaluation is publicly available in the https://github.com/QSong-github/MVCBench. The repository also provides scripts for dataset downloading and processing, the main framework implementation, plotting utilities for reproducing the figures, and all experimental results.

## ACKNOWLEDGEMENTS

B. Z. was supported by the University of Macau [Grant Number MYRG-GRG2024-00205-FST]; and the Science and Technology Development Fund, Macau SAR [Grant Number 0028/2023/RIA1]

## TABLES

**Table 1: Benchmarking datasets for model performance evaluation**.

**Table 2: Parameter specifications of benchmarked foundation models**.

## SUPPLEMENTARY INFORMATION

**Supplementary Fig. 1 Data scale, composition, and technical variation analysis of the benchmarking dataset**.

**Supplementary Fig. 2 Extended performance metrics and cross-plate generalization analysis of drug molecular representations**.

**Supplementary Fig. 3 Validation of drug molecular representation performance on the BBBC036_Image dataset.**

**Supplementary Fig. 4 Extended validation and generalization analysis of gene representation methods**.

**Supplementary Fig. 5 Impact of multimodal fusion strategies on gene expression prediction and validation on BBBC047 (left) and BBBC036 (right) datasets**.

**Supplementary Fig. 6 Validation of modality orthogonality**.

**Supplementary Note 1: Quantification of Experimental Heterogeneity and Transfer Difficulty.**

**Supplementary Note 2: Model Architectures and Implementation Details**.

## REFERENCES

1. Tang, L. The virtual cell. Nat. Methods 22, 2493–2493 (2025).

2. Way, G. P. et al. Morphology and gene expression profiling provide complementary information for mapping cell state. Cell Syst. 13, 911-923.e9 (2022).

3. Bray, M.-A. et al. Cell Painting, a high-content image-based assay for morphological profiling using multiplexed fluorescent dyes. Nat. Protoc. 11, 1757–1774 (2016).

4. Heumos, L. et al. Pertpy: an end-to-end framework for perturbation analysis. Preprint at 10.1101/2024.08.04.606516 (2024).

5. Ergen, C. et al. Scvi-hub: an actionable repository for model-driven single cell analysis. Preprint at 10.1101/2024.03.01.582887 (2024).

6. Cui, H. et al. Towards multimodal foundation models in molecular cell biology. Nature 640, 623–633 (2025).

7. Bunne, C. et al. How to build the virtual cell with artificial intelligence: Priorities and opportunities. Cell 187, 7045–7063 (2024).

8. Johnson, J. A. I. et al. Human interpretable grammar encodes multicellular systems biology models to democratize virtual cell laboratories. Cell 188, 4711-4733.e37 (2025).

9. Qian, L., Dong, Z. & Guo, T. Grow AI virtual cells: three data pillars and closed-loop learning. Cell Res. 35, 319–321 (2025).

10. Haghighi, M., Caicedo, J. C., Cimini, B. A., Carpenter, A. E. & Singh, S. High-dimensional gene expression and morphology profiles of cells across 28,000 genetic and chemical perturbations. Nat. Methods 19, 1550–1557 (2022).

11. Chandrasekaran, S. N. et al. JUMP Cell Painting dataset: morphological impact of 136,000 chemical and genetic perturbations. Preprint at 10.1101/2023.03.23.534023 (2023).

12. Peidli, S. et al. scPerturb: harmonized single-cell perturbation data. Nat. Methods 21, 531–540 (2024).

13. Subramanian, A. et al. A Next Generation Connectivity Map: L1000 Platform and the First 1,000,000 Profiles. Cell 171, 1437-1452.e17 (2017).

14. Verma, S. et al. Generating Joint Transcriptomic and Morphological Responses to Drug Perturbations via Rectified Flow. 10.64898/2026.02.02.703189 (2026).

15. Li, C. et al. Benchmarking AI Models for In Silico Gene Perturbation of Cells. Preprint at 10.1101/2024.12.20.629581 (2024).

16. Hu, Y. et al. Benchmarking algorithms for single-cell multi-omics prediction and integration. Nat. Methods 21, 2182–2194 (2024).

17. Partin, A. et al. Benchmarking community drug response prediction models: datasets, models, tools, and metrics for cross-dataset generalization analysis. Preprint at 10.48550/arXiv.2503.14356 (2025).

18. Cui, H. et al. scGPT: toward building a foundation model for single-cell multi-omics using generative AI. Nat. Methods 21, 1470–1480 (2024).

19. Rogers, D. & Hahn, M. Extended-Connectivity Fingerprints. J. Chem. Inf. Model. 50, 742–754 (2010).

20. Bosch, I. et al. Benchmarking knowledge graph embedding models for the prediction of oligogenic combinations. Brief. Bioinform. 27, bbaf712 (2026).

21. Wu, Z. et al. A Comprehensive Survey on Graph Neural Networks. IEEE Trans. Neural Netw. Learn. Syst. 32, 4–24 (2021).

22. Heid, E. et al. Chemprop: A Machine Learning Package for Chemical Property Prediction. J. Chem. Inf. Model. 64, 9–17 (2024).

23. Li, H. et al. A knowledge-guided pre-training framework for improving molecular representation learning. Nat. Commun. 14, 7568 (2023).

24. Seal, S. et al. Integrating cell morphology with gene expression and chemical structure to aid mitochondrial toxicity detection. Commun. Biol. 5, 858 (2022).

25. He, S. et al. Squidiff: predicting cellular development and responses to perturbations using a diffusion model. Nat. Methods. 10.1038/s41592-025-02877-y (2025).

26. Roohani, Y., Huang, K. & Leskovec, J. Predicting transcriptional outcomes of novel multigene perturbations with GEARS. Nat. Biotechnol. 42, 927–935 (2024).

27. Wang, X. et al. Multimodal pre-training models of molecular representation for drug discovery. Natl. Sci. Rev. 13, nwaf495 (2026).

28. Lejal, V., Rouquié, D. & Taboureau, O. Cell morphology and gene expression: tracking changes and complementarity across time and cell lines. Preprint at 10.1101/2024.08.30.610494 (2024).

29. Cerisier, N., Dafniet, B., Badel, A. & Taboureau, O. Linking chemicals, genes and morphological perturbations to diseases. Toxicol. Appl. Pharmacol. 461, 116407 (2023).

30. Broad Institute. Expanded CMap LINCS Resource 2020. https://clue.io/data/CMap2020#LINCS2020.

31. Zhang, J. et al. Tahoe-100M: A Giga-Scale Single-Cell Perturbation Atlas for Context-Dependent Gene Function and Cellular Modeling. Preprint at 10.1101/2025.02.20.639398 (2025).

32. Xiang, L. et al. High-throughput profiling of chemical-induced gene expression across 93,644 perturbations. Nat. Methods. 10.1038/s41592-025-02781-5 (2025).

33. Chandrasekaran, S. N. et al. JUMP Cell Painting dataset: morphological impact of 136,000 chemical and genetic perturbations. Preprint at 10.1101/2023.03.23.534023 (2023).

34. Wang, Y., Wang, J., Cao, Z. & Barati Farimani, A. Molecular contrastive learning of representations via graph neural networks. Nat. Mach. Intell. 4, 279–287 (2022).

35. Liu, G. et al. Learning Molecular Representation in a Cell. Preprint at http://arxiv.org/abs/2406.12056 (2024).

36. Ahmad, W., Simon, E., Chithrananda, S., Grand, G. & Ramsundar, B. ChemBERTa-2: Towards Chemical Foundation Models. Preprint at 10.48550/arXiv.2209.01712 (2022).

37. Xia, J. et al. Mole-BERT: Rethinking Pre-training Graph Neural Networks for Molecules. in The Eleventh International Conference on Learning Representations (2023). doi:10.26434/chemrxiv-2023-dngg4.

38. Edwards, C. et al. Translation between Molecules and Natural Language. in Proceedings of the 2022 Conference on Empirical Methods in Natural Language Processing (arXiv, 2022). doi:10.48550/arXiv.2204.11817.

39. Zhou, G. et al. Uni-Mol: A Universal 3D Molecular Representation Learning Framework. in The Eleventh International Conference on Learning Representations (2023).

40. Ji, X. et al. Uni-Mol2: Exploring Molecular Pretraining Model at Scale. Preprint at 10.48550/arXiv.2406.14969 (2024).

41. Wang, L. et al. Conformational Space Profiling Enhances Generic Molecular Representation for AI-Powered Ligand-Based Drug Discovery. Adv. Sci. 11, 2403998 (2024).

42. Wang, L. et al. Directed Chemical Evolution via Navigating Molecular Encoding Space. Preprint at 10.1101/2025.03.18.643899 (2025).

43. Theodoris, C. V. et al. Transfer learning enables predictions in network biology. Nature 618, 616–624 (2023).

44. Yang, F. et al. scBERT as a large-scale pretrained deep language model for cell type annotation of single-cell RNA-seq data. Nat. Mach. Intell. 4, 852–866 (2022).

45. Zhao, S., Zhang, J. & Nie, Z. Large-Scale Cell Representation Learning via Divide-and-Conquer Contrastive Learning. Preprint at 10.48550/arXiv.2306.04371 (2023).

46. Heimberg, G. et al. A cell atlas foundation model for scalable search of similar human cells. Nature 638, 1085–1094 (2025).

47. Hao, M. et al. Large-scale foundation model on single-cell transcriptomics. Nat. Methods 21, 1481–1491 (2024).

48. Rosen, Y. et al. Universal Cell Embeddings: A Foundation Model for Cell Biology. Preprint at 10.1101/2023.11.28.568918 (2023).

49. Wen, H. et al. CellPLM: Pre-training of Cell Language Model Beyond Single Cells. in Proceedings of the Twelfth International Conference on Learning Representations (ICLR 2024). doi:10.1101/2023.10.03.560734.

50. Adduri, A. K. et al. Predicting cellular responses to perturbation across diverse contexts with State. 10.1101/2025.06.26.661135 (2026).

51. Shen, H. et al. Generative pretraining from large-scale transcriptomes: Implications for single-cell deciphering and clinical translation. Preprint at 10.1101/2022.01.31.478596 (2022).

52. Levine, D. et al. Cell2Sentence: Teaching Large Language Models the Language of Biology. in Proceedings of the 41st International Conference on Machine Learning (Bioinformatics, 2023). doi:10.1101/2023.09.11.557287.

53. Viñas Torné, R. et al. Systema: a framework for evaluating genetic perturbation response prediction beyond systematic variation. Nat. Biotechnol. 10.1038/s41587-025-02777-8 (2025).

54. Li, B. et al. PhenoProfiler: Advancing Phenotypic Learning for Image-based Drug Discovery. Nat. Commun. (2025).

55. Wang, S., et al. PhenoScreen: A Dual-Space Contrastive Learning Framework-based Phenotypic Screening Method by Linking Chemical Perturbations to Cellular Morphology. bioRxiv 10.1101/2024.10.23.619752 (2024).

56. Heumos, L. et al. Best practices for single-cell analysis across modalities. Nat. Rev. Genet. 24, 550–572 (2023).

57. Vieth, B., Parekh, S., Ziegenhain, C., Enard, W. & Hellmann, I. A systematic evaluation of single cell RNA-seq analysis pipelines. Nat. Commun. 10, 4667 (2019).

58. Chen, S., Zhang, Y. & Yang, Q. Multi-Task Learning in Natural Language Processing: An Overview. ACM Comput. Surv. 56, 1–32 (2024).

59. Kingma, D. P. & Ba, J. Adam: A Method for Stochastic Optimization. Preprint at 10.48550/arXiv.1412.6980 (2017).

