## Supplemental Data 1 for "MVCBench: A Multimodal Benchmark for Drug-induced Virtual Cell Phenotypes"

Supplementary Figures ---------------- 3-7

Supplementary Notes ---------------- 8-10

Supplementary References ---------------- 11

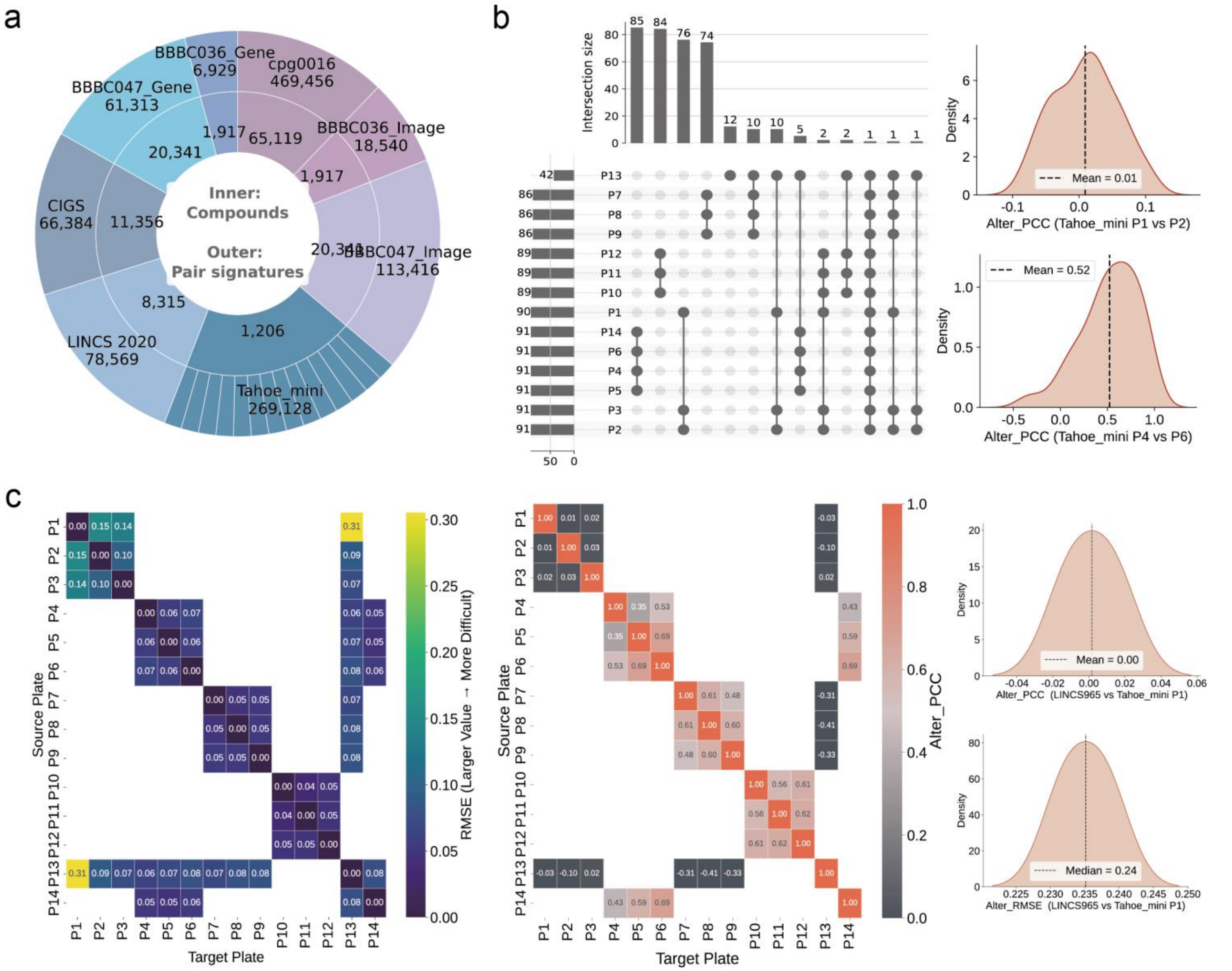

**Supplementary Fig. 1 Data scale, composition, and technical variation analysis of the benchmarking dataset**. **a**, Sunburst chart illustrating the scale of the benchmark datasets. The inner ring represents the number of unique compounds, and the outer ring denotes the count of paired drug-induced data across transcriptomic and morphological modalities. **b**, Compound overlap and reproducibility analysis in the Tahoe_mini dataset. Left: Upset plot displaying the intersection of compounds tested across different plates. Right: Density plots of replicate correlations (Alter_PCC) revealing distinct batch consistency; for instance, P1 vs. P2 shows near-zero correlation (indicating high technical noise/batch effects), whereas P4 vs. P6 shows high correlation (indicating consistent biological signals). **c**, Quantification of transfer difficulty across plates and datasets. Left/Middle: Cross-plate similarity matrices based on RMSE and PCC for Tahoe_mini. Plates P1 and P13 exhibit high error and low correlation with other batches, identifying them as challenging outlier scenarios used for leave-plate-out generalization evaluation. Right: Distribution of correlation (Alter_PCC) and error (Alter_RMSE) when transferring from LINCS2020 (965 features) to Tahoe_mini, quantifying the fundamental domain shift between two transcriptomic datasets.

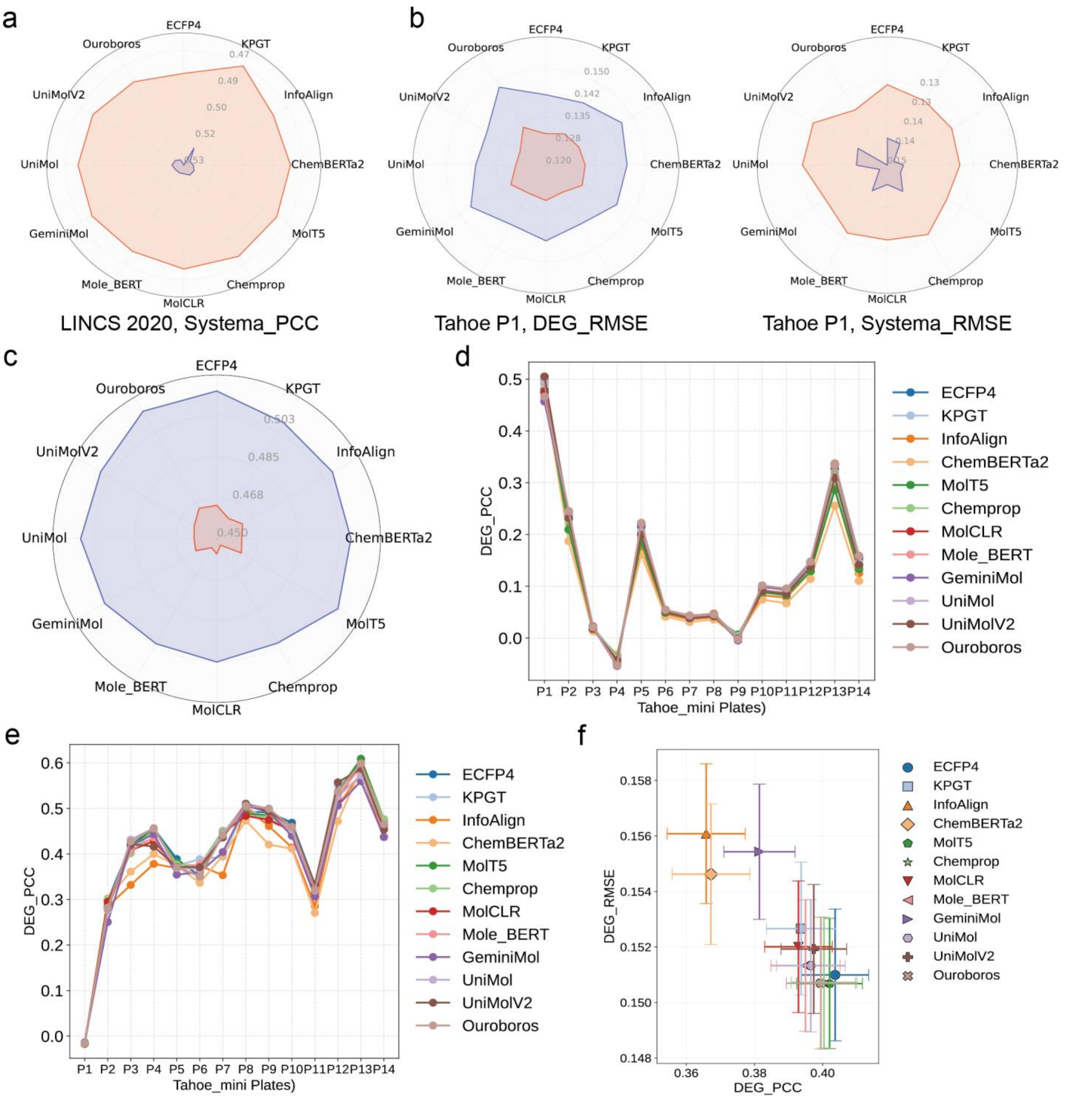

**Supplementary Fig. 2 Extended performance metrics and cross-plate generalization analysis of drug molecular representations. a**, Radar chart showing the Systema_PCC performance of 12 drug molecular representation methods on the LINCS2020 dataset. **b**, Extended evaluation on the Tahoe_mini P1 dataset. Radar charts display DEG_RMSE (Left) and Systema_RMSE (Right). **c**, Extended evaluation of DEG_RMSE metric on the LINCS2020 dataset under leave-cell-line-out setting. **d–e**, Cross-plate generalization performance on the Tahoe_mini dataset. The plots illustrate the predictive fidelity (DEG_PCC) of models trained on challenging outlier plates P1 (d) and P13 (e) and evaluated across all other plates to assess robustness to technical batch effects. **f**, Scatter plot summarizing the overall trade-off between predictive accuracy (DEG_PCC) and error (DEG_RMSE) for all evaluated molecular representation methods.

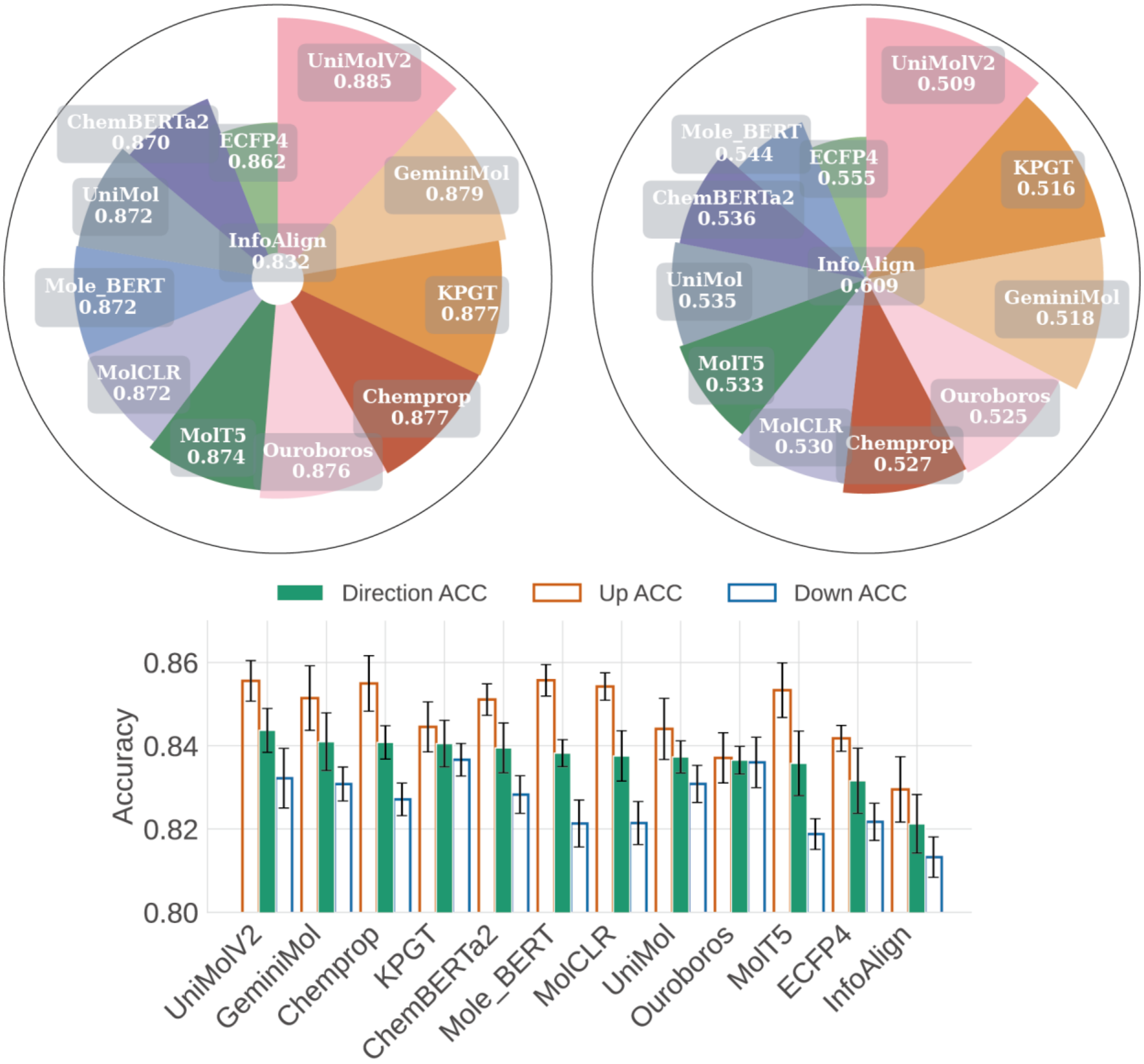

**Supplementary Fig. 3 Validation of drug molecular representation performance on the BBBC036_Image dataset.** Comparative analysis of predictive fidelity (DEG_PCC) across varying molecular encoders on the BBBC036_Image dataset.

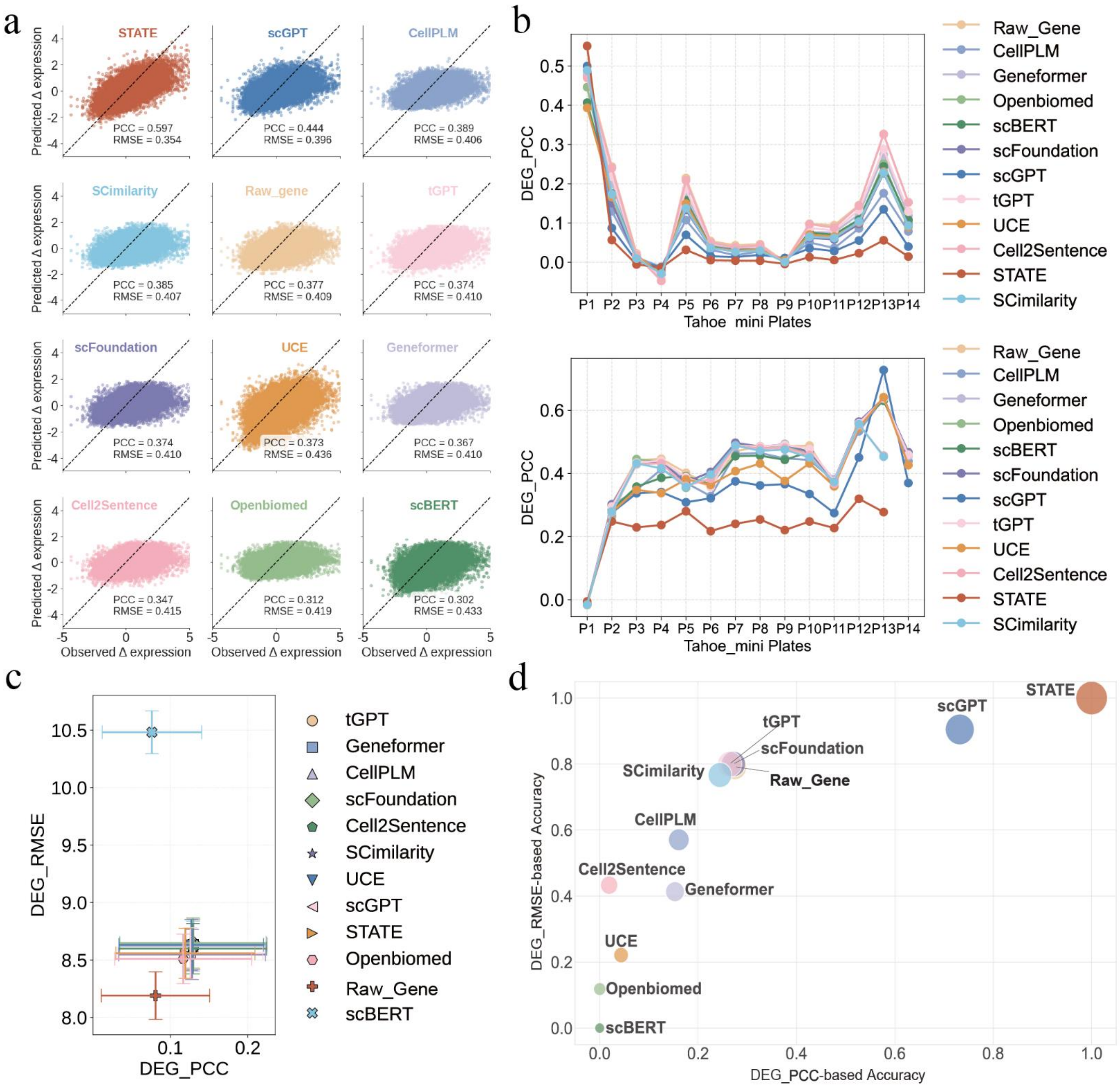

**Supplementary Fig. 4 Extended validation and generalization analysis of gene representation methods. a**, Predictive validation on the high-expressed genes subset of Tahoe_mini P1 dataset. **b**, Plate-specific generalization performance on the Tahoe_mini dataset. Line plots display DEG_PCC across 14 target plates when models are trained specifically on the outlier plate P13 (top) or P1 (bottom). **c**, Cross-dataset zero-shot transfer performance (Tahoe_mini → LINCS2020). Scatter plot of DEG_RMSE versus DEG_PCC showing the severe performance collapse when models trained on Tahoe_mini data are transferred to predict drug-induced responses in LINCS2020 dataset. **d**, multi-dimensional performance landscape of gene representations. Bubble plot illustrating the relative positioning of different representation methods based on their normalized average DEG_PCC (x-axis) and DEG_RMSE (y-axis). The spatial distribution highlights the trade-off between predictive correlation and error magnitude, with methods in the upper-right quadrant, such as STATE and scGPT, demonstrating superior robustness across multimodal benchmarking tasks.

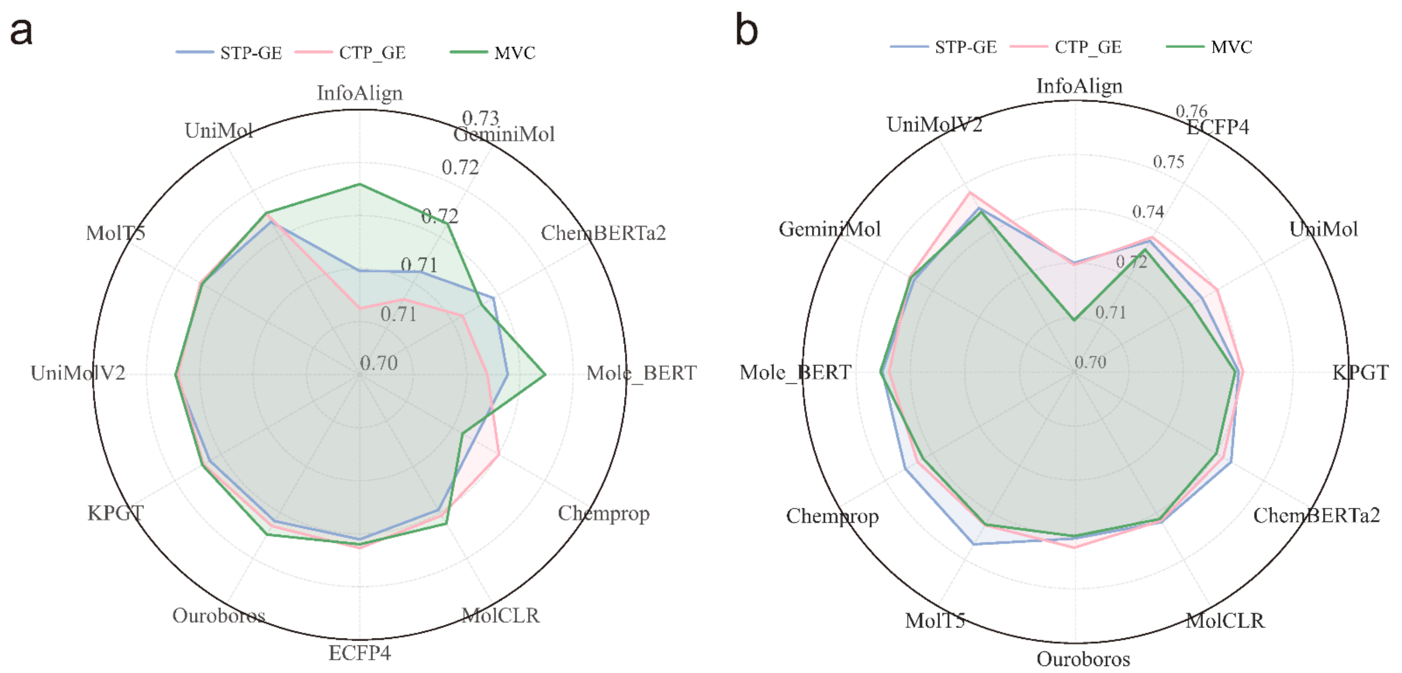

**Supplementary Fig. 5 Impact of multimodal fusion strategies on gene expression prediction and validation on BBBC047 (left) and BBBC036 (right) datasets. a**, Comparative analysis of gene expression prediction performance (DEG_PCC) under three modeling paradigms: single-task paradigm (STP-GE), in which the model receives drug structure and pre-treatment gene expression only and is optimized with a single-task objective; cross-modal assisted paradigm (CTP-GE), which incorporates pre-treatment morphological profiles as auxiliary input while retaining a single-task supervision on gene expression; and multimodal virtual cell (MVC), where both modalities are provided as inputs and jointly optimized under a multi-task objective. **b**, Cross-dataset validation on the BBBC036 benchmark. Consistent performance gains observed under CTP-GE and MVC demonstrate that improvements arising from cross-modal contextualization and joint optimization are reproducible across independent datasets, supporting the robustness and generalizability of the MVC framework beyond the primary LINCS2020/Tahoe_mini benchmark.

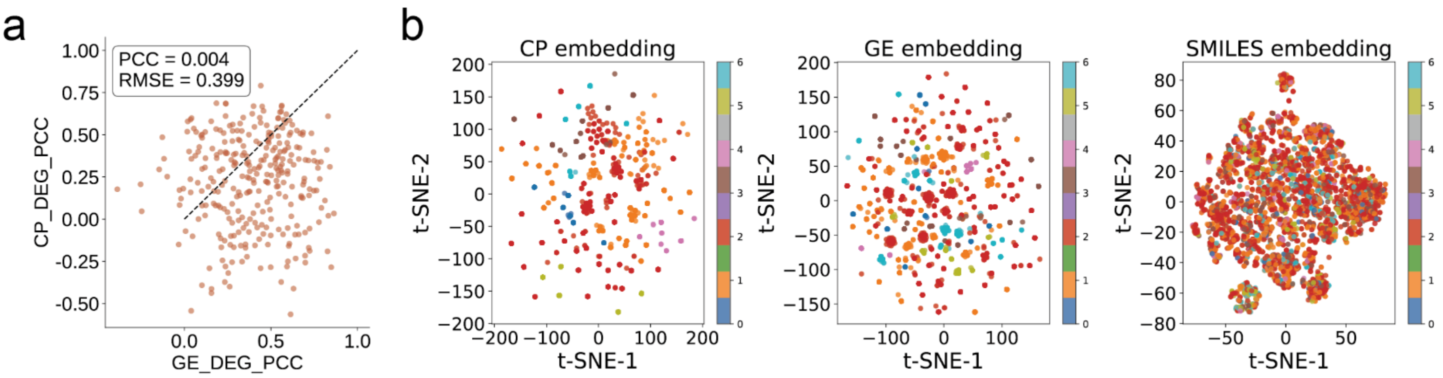

**Supplementary Fig. 6 Validation of modality orthogonality. a**, Scatter plot comparing the predictive performance of gene expression versus morphology on the BBBC036 dataset. The near-zero correlation confirms that these two modalities provide independent biological information. **b**, t-SNE visualization of the joint latent space. The clear separation between drug (SMILES), transcriptomic (GE), and morphological (CP) embeddings illustrates that these modalities occupy distinct, non-overlapping data manifolds.

### SUPPLEMENTARY NOTES

#### Note 1: Quantification of Experimental Heterogeneity and Transfer Difficulty

##### (1) Quantifying domain shift with Alter_PCC

To systematically quantify the magnitude of distributional shifts between experimental domains (e.g., between plates or independent datasets), we introduced the Inter-Domain Consistency metric (Alter_PCC). This metric evaluates the reproducibility of ground-truth drug-induced signatures across domains prior to model evaluation. The calculation proceeds in three steps: 1) Differential calculation**:** For each domain, we compute the differential expression profile $\delta = x_{treated} - x_{control}$ for every sample. 2) Consensus aggregation: We mitigate replicate noise by computing a consensus signature $v_d$ for each drug $d$. This is obtained by averaging the differential profiles $\delta$ across all $N$ biological replicates within the domain:

$$v_d = \frac{1}{N}\sum_{i=1}^{N} \delta_i$$

3) Metric computation**:** We identify the set of $K$ compounds shared between the source domain ($D_{src}$) and target domain ($D_{tgt}$). The consensus signatures for these shared compounds are stacked to form aligned matrices. Alter_PCC is defined as the Pearson correlation coefficient between the flattened vectors of these two matrices:

$$Alter\_PCC = PCC(\text{flatten}(V_{src}),\text{flatten}(V_{tgt}))$$

An Alter_PCC approaching zero indicates a lack of biological consistency between domains, quantifying the intrinsic difficulty of the transfer task.

##### (2) Inter-plate transfer difficulty and batch effects

To interpret the generalization barriers observed in cross-plate evaluations, we quantified the impact of batch effects on transferability. Standardized drug-induced transcriptional signatures were extracted for compounds shared across all 14 plates, and inter-plate coverage and response similarity matrices were computed (**Supplementary Fig. 1b, c**). This analysis revealed generally low inter-plate correlations and many plate pairs with insufficient shared compounds, highlighting substantial experimental heterogeneity. Importantly, we observed a strong positive correlation between distributional similarity and generalization potential, indicating that performance degradation reported in the main text primarily arises from intrinsic data incompatibility caused by technical batch effects rather than algorithmic limitations.

##### (3) Cross-dataset domain shift

To assess transferability between LINCS2020[1,2] and Tahoe_mini[3], we measured inter-domain consistency using Alter_PCC, which quantifies the correlation of ground-truth drug response signatures for shared compounds across domains. Our analysis revealed a severe distributional shift (Alter_PCC = 0.00, Alter_RMSE = 0.24; **Supplementary Fig. 1d**), demonstrating near-total lack of biological concordance between datasets. This intrinsic incompatibility explains the sharp loss of predictive performance observed in cross-dataset evaluations.

#### Note 2: Model Architectures and Implementation Details

We benchmarked 24 representation methods spanning drug molecular and gene expression modalities. All models were deployed using official repositories and publicly released pre-trained weights, with parameters frozen during downstream evaluation. Architectural specifications and pretraining configurations are detailed below.

##### (1) Drug molecular representation methods

We benchmarked 12 methods categorized into rule-based, graph-based, Transformer-based, and geometry-aware architectures. The rule-based baseline employed ECFP4[4] fingerprints (radius = 2, bit length = 2048) generated via RDKit, which were utilized directly as raw binary vectors without additional scaling or normalization. GNN-based models included Chemprop[5], MolCLR[6], and InfoAlign[7], which operate directly on molecular graphs. Chemprop leverages a message-passing neural network (MPNN) to aggregate atomic features; we used the official pretrained checkpoint (*example_model_v2_regression_mol.ckpt*) with the penultimate feed-forward layer as output. MolCLR applies contrastive learning with graph augmentations (e.g., subgraph deletion) to enhance robustness, while InfoAlign maximizes mutual information between local and global graph views. Both were deployed using official repositories and pretrained weights.

Transformer-based models, specifically ChemBERTa2[8], Mole-BERT[9], MolT5[10], and KPGT[11], encode SMILES or graph structures leveraging self-attention to capture long-range dependencies. ChemBERTa2, built on RoBERTa[12], was applied using the *ChemBERTa-77M-MLM* checkpoint. Mole-BERT integrates contrastive learning into graph Transformers[13], while MolT5 adopts a text-to-text generative pretraining paradigm. KPGT incorporates external knowledge graphs to jointly model molecular structure and domain knowledge. All models were deployed following official guidelines with publicly released weights. Finally, geometry-aware models, including UniMol[14], UniMolV2[15], GeminiMol[16], and Ouroboros[17], extend graph-based representations by incorporating 3D coordinates. UniMol and UniMolV2 employ large-scale self-supervised pretraining on molecular conformations; we used the 84M and 310M parameter versions, respectively. GeminiMol adopts a dual-branch architecture to jointly encode 2D and 3D representations, whereas Ouroboros leverages iterative generation and contrastive learning to stabilize geometric embeddings. Both were implemented using official repositories and pretrained checkpoints. For all drug representation methods, we utilized the pre-trained checkpoints to extract molecular features without fine-tuning the underlying encoders on our benchmark tasks.

**(2) Gene representation methods**

We benchmarked 12 gene representation methods for drug-induced gene expression prediction, comprising a Raw_Gene baseline and 11 deep learning models spanning three major architectural paradigms: encoder-only (Geneformer[18], scBERT[19], OpenBioMed[20], SCimilarity[21]), encoder–decoder (scFoundation[22], scGPT[23], UCE[24], CellPLM[25], STATE[26]), and decoder-only (tGPT[27], Cell2Sentence[28]). For the Raw_Gene baseline, the pre-treatment gene expression profile was used directly as the cellular embedding without passing through any pre-trained foundation model encoder, serving as a raw feature input to the downstream predictor.

Encoder-only models focus on learning efficient representations from single-cell data, typically adopting BERT-like architectures with self-attention to capture complex gene–gene dependencies. Geneformer leverages large-scale pretraining on single-cell transcriptomes to learn a universal “language” of gene expression, enabling robust inference of cell type, state, and perturbation effects. scBERT treats gene expression profiles as sentences and genes as tokens, applying masked language modeling to derive informative embeddings. OpenBioMed provides a broader biomedical pretraining framework, with its single-cell models (e.g., CellLM) designed to extract generalizable biological knowledge from large-scale datasets. SCimilarity is a deep metric-learning framework designed to learn a unified and interpretable cell representation by jointly optimizing unsupervised representation learning and a supervised triplet loss function. It is primarily used for scalable search and quantification of similarity between cell profiles.

Encoder-decoder models extend beyond representation learning to generative tasks, leveraging sequence-to-sequence architectures inspired by machine translation. scFoundation aims to serve as a universal backbone for single-cell analysis, integrating diverse data types to enhance cross-modal and cross-species generalization. scGPT adapts generative pretraining to model single-cell data as sequences, supporting tasks such as cell type classification and perturbation effect prediction. UCE (Universal Cell Embeddings) employs contrastive learning within an encoder–decoder framework to produce robust embeddings while modeling perturbation responses. CellPLM conceptualizes single-cell data as a biological language, using encoder-decoder pretraining to capture the semantic complexity of cellular states. STATE employs an encoder-decoder architecture where a dense, bidirectional Transformer encoder predicts log-normalized gene expression, coupled with a specialized multi-layer perceptron (MLP) decoder to map embeddings back to the expression space.

Finally, decoder-only models, exemplified by tGPT, adopt an autoregressive paradigm similar to GPT. By modeling single-cell transcriptomes as ordered sequences, tGPT predicts gene expression in a self-regressive manner, making it particularly suited for capturing temporal or spatial dependencies and simulating gene regulatory dynamics. Cell2Sentence (C2S) is a method designed to directly adapt large language models (LLMs) to single-cell transcriptomics. It converts single-cell gene expression data into "cell sentences" and then fine-tunes existing LLM architectures, such as GPT-2, which are typically decoder-only models, for tasks like cell generation and perturbation prediction.

#### References

1. Subramanian, A. *et al.* A Next Generation Connectivity Map: L1000 Platform and the First 1,000,000 Profiles. *Cell* **171**, 1437-1452.e17 (2017).
2. Broad Institute. Expanded CMap LINCS Resource 2020. https://clue.io/data/CMap2020#LINCS2020.
3. Zhang, J. *et al.* Tahoe-100M: A Giga-Scale Single-Cell Perturbation Atlas for Context-Dependent Gene Function and Cellular Modeling. Preprint at https://doi.org/10.1101/2025.02.20.639398 (2025).
4. Rogers, D. & Hahn, M. Extended-Connectivity Fingerprints.
5. Heid, E. *et al.* Chemprop: A Machine Learning Package for Chemical Property Prediction. *J. Chem. Inf. Model.* **64**, 9–17 (2024).
6. Wang, Y., Wang, J., Cao, Z. & Barati Farimani, A. Molecular contrastive learning of representations via graph neural networks. *Nat. Mach. Intell.* **4**, 279–287 (2022).
7. Liu, G. *et al.* Learning Molecular Representation in a Cell. Preprint at http://arxiv.org/abs/2406.12056 (2024).
8. Ahmad, W., Simon, E., Chithrananda, S., Grand, G. & Ramsundar, B. ChemBERTa-2: Towards Chemical Foundation Models. Preprint at https://doi.org/10.48550/arXiv.2209.01712 (2022).
9. Xia, J. *et al.* Mole-BERT: Rethinking Pre-training Graph Neural Networks for Molecules. in *The Eleventh International Conference on Learning Representations* (2023). doi:10.26434/chemrxiv-2023-dngg4.
10. Edwards, C. *et al.* Translation between Molecules and Natural Language. in *Proceedings of the 2022 Conference on Empirical Methods in Natural Language Processing* (arXiv, 2022). doi:10.48550/arXiv.2204.11817.
11. Li, H. *et al.* A knowledge-guided pre-training framework for improving molecular representation learning. *Nat. Commun.* **14**, 7568 (2023).
12. Liu, Y. *et al.* RoBERTa: A Robustly Optimized BERT Pretraining Approach. Preprint at https://doi.org/10.48550/arXiv.1907.11692 (2019).
13. Shehzad, A. *et al.* Graph Transformers: A Survey. *IEEE Trans. Neural Netw. Learn. Syst.* 1–20 (2026) doi:10.1109/TNNLS.2025.3646122.
14. Zhou, G. *et al.* Uni-Mol: A Universal 3D Molecular Representation Learning Framework. in *The Eleventh International Conference on Learning Representations* (2023).
15. Ji, X. *et al.* Uni-Mol2: Exploring Molecular Pretraining Model at Scale. Preprint at https://doi.org/10.48550/arXiv.2406.14969 (2024).
16. Wang, L. *et al.* Conformational Space Profiling Enhances Generic Molecular Representation for AI-Powered Ligand-Based Drug Discovery. *Adv. Sci.* **11**, 2403998 (2024).
17. Wang, L. *et al.* Directed Chemical Evolution via Navigating Molecular Encoding Space. Preprint at https://doi.org/10.1101/2025.03.18.643899 (2025).
18. Theodoris, C. V. *et al.* Transfer learning enables predictions in network biology. *Nature* **618**, 616–624 (2023).
19. Yang, F. *et al.* scBERT as a large-scale pretrained deep language model for cell type annotation of single-cell RNA-seq data. *Nat. Mach. Intell.* **4**, 852–866 (2022).
20. Zhao, S., Zhang, J. & Nie, Z. Large-Scale Cell Representation Learning via Divide-and-Conquer Contrastive Learning. Preprint at https://doi.org/10.48550/arXiv.2306.04371 (2023).

21. Heimberg, G. *et al.* A cell atlas foundation model for scalable search of similar human cells. *Nature* **638**, 1085–1094 (2025).

22. Hao, M. *et al.* Large-scale foundation model on single-cell transcriptomics. *Nat. Methods* **21**, 1481–1491 (2024).

23. Cui, H. *et al.* scGPT: toward building a foundation model for single-cell multi-omics using generative AI. *Nat. Methods* **21**, 1470–1480 (2024).

24. Rosen, Y. *et al.* Universal Cell Embeddings: A Foundation Model for Cell Biology. Preprint at https://doi.org/10.1101/2023.11.28.568918 (2023).

25. Wen, H. *et al.* CellPLM: Pre-training of Cell Language Model Beyond Single Cells. in *Proceedings of the Twelfth International Conference on Learning Representations (ICLR 2024).* doi:10.1101/2023.10.03.560734.

26. Adduri, A. K. *et al.* Predicting cellular responses to perturbation across diverse contexts with State.

27. Shen, H. *et al.* Generative pretraining from large-scale transcriptomes: Implications for single-cell deciphering and clinical translation. Preprint at https://doi.org/10.1101/2022.01.31.478596 (2022).

28. Levine, D. *et al.* Cell2Sentence: Teaching Large Language Models the Language of Biology. in *Proceedings of the 41st International Conference on Machine Learning* (Bioinformatics, 2023). doi:10.1101/2023.09.11.557287.
